# Mitochondrial double-stranded RNAs as a pivotal mediator in the pathogenesis of Sjögren’s syndrome

**DOI:** 10.1101/2021.09.13.459934

**Authors:** Jimin Yoon, Minseok Lee, Ahsan Ausaf Ali, Ye Rim Oh, Yong Seok Choi, Sujin Kim, Namseok Lee, Se Gwang Jang, Seung-Ki Kwok, Joon Young Hyon, Seunghee Cha, Yun Jong Lee, Sung Gap Im, Yoosik Kim

## Abstract

**Objective:** Sjögren’s syndrome (SS) is a systemic autoimmune disease that targets the exocrine glands, resulting in impaired saliva and tear secretion. To date, type I interferons (I-IFNs) are increasingly recognized as pivotal mediators in SS, but their endogenous drivers have not been elucidated. This study investigates the role of mitochondrial double-stranded RNAs (mt-dsRNAs) in regulating I-IFN response in SS.

**Methods:** Saliva and tear from SS patients and controls (n=73 for saliva and n=16 for tear), the salivary glands of the SS-prone-non-obese-diabetic mouse, and primary human salivary glandular cells were screened for mt-dsRNAs by RT-qPCR. The human salivary cell line (NS-SV-AC) grown as three-dimensional spheroids were subject to dsRNA stress to measure mt-dsRNA induction and recapitulation of SS glandular inflammatory features. Acetylcholine, SS-IgG, upadacitinib (JAK1 inhibitor), or 2-C′-methyladenosine (mitochondrial transcription inhibitor) were applied to characterize the roles of mt-dsRNAs. To identify endogenous dsRNA-sensor and confirm the mitochondrial origin of cytoplasmic dsRNAs, the immunoprecipitation of dsRNAs was performed.

**Results:** mt-dsRNAs were elevated in the SS specimens with salivary ND5 and tear CYTB1 being statistically associated with secretory dysfunction/inflammation and corneal/conjunctival damage, respectively. Stimulation of the spheroids with dsRNA stress of poly I:C induced mt-dsRNAs, p-PKR, and I-IFNS via the JAK1/STAT pathway whereas the inhibition of mt-RNA synthesis or JAK1 attenuated the glandular signature. The inhibitory effect of acetylcholine on mt-dsRNAs and I-IFNS induction was reversed by SS-IgG.

**Conclusion:** mt-dsRNAs amplify the impact of dsRNA stress on SS glandular signatures *in vitro*, potentially propagating a pseudo-viral signal in the SS target tissue.

**Summary:** Mitochondrial double-stranded RNA levels were elevated in the tear and saliva of SS patients, which was associated with secretory dysfunction and tissue inflammation. These RNAs amplified type I interferon signature as well as glandular phenotypes reported in SS. Inhibitors of mitochondrial RNA transcription or JAK1 in salivary gland acinar cell spheroids attenuated the mitochondrial RNA-mediated changes.

## INTRODUCTION

Sjögren’s syndrome (SS) is a chronic disorder characterized by lymphocytic infiltration in the exocrine glands and B-cell hyperactivity with autoantibody production, resulting in oral and eye dryness. Similar to most autoimmune disorders, the etiology of SS is not yet fully understood. It can be challenging to diagnose patients with SS due to heterogeneity of disease manifestations and the lack of disease-specific biomarkers for the condition [1, 2]. A delay in timely diagnosis and intervention can adversely affect patients’ quality of life due to severe hyposalivation, glandular B-cell lymphoma, and/or extraglandular organ involvement such as interstitial lung disease [3].

One of the most notable features of SS is the type I interferon (I-IFN) signature detected in the minor salivary gland lip biopsy specimens, CD14+ blood monocytes, plasmacytoid dendritic cells, and PBMC [4, 5]. I-IFNs, which are critical molecules in host defense against viral infections, can be produced in response to stimulation of pattern recognition receptors (PRRs). In particular, toll-like receptors (TLRs), protein kinase R (PKR), and melanoma differentiation-associated gene 5 (MDA5) are known to be stimulated by double-stranded RNAs (dsRNAs) that are often originated from the viral genome [6, 7]. The upregulation of TLRs is accompanied by increased levels of IFN-stimulated genes (ISGs) and proinflammatory cytokines in the salivary glands of SS [8-10]. The above-mentioned findings suggest that the dysregulated dsRNAs play a role in the pathogenesis of SS [7].

Furthermore, recent evidence indicates that dsRNAs endogenously produced in host cells in the absence of infection can activate PRRs [11, 12]. Among them, those originated from the bidirectional transcription of the mitochondrial genome (mt-dsRNAs) have received great attention due to their pathological implications in the context of alcohol-induced liver injury, Huntington’s disease, and osteoarthritis [13-15]. However, potential roles of mt-dsRNAs in SS or other autoimmune conditions have never been investigated to our knowledge. When mt-dsRNAs are aberrantly present in the cytosol, they can be recognized by PKR and/or MDA5, resulting in downstream signal cascades for immune responses [11, 16]. Considering the excess level of oxidative stress and mitochondrial dysfunction were reported in the salivary glands of SS [17-20], we hypothesize that the generation of mt-dsRNAs, potentially from altered mitochondrial bioenergetics in the inflamed target tissues, further stimulates PRRs and amplifies I-IFN-mediated inflammatory responses in salivary glands of SS.

The aberrant immune responses in the target glandular tissues of SS are known to alter expression and distribution of water channel proteins, such as aquaporin 5 (AQP5) and/or tight junction complex (TJC) proteins [21, 22]. The presence of autoantibodies against acinar cell surface receptor, muscarinic type 3 receptor (M3R), is also known to suppress acetylcholine (Ach)-mediated secretory function in the salivary glands of SS [23]. In addition to acinar cell death caused by infiltrated autoreactive cytotoxic T cells, soluble factors, such as cytokines or anti-M3R autoantibodies, are known to inhibit the expression of AQP5 and TCJ proteins and Ach-stimulation of M3R, respectively, resulting in secretory dysfunction. These factors may explain why SS patients lose secretory function even in the absence of cytotoxic T cells in the target salivary glands.

In our current study, we investigated the roles of mt-dsRNA in SS for the first time, with a special focus on their impact on the glandular autoimmune signature reported in the disease. We screened the presence of mt-dsRNAs in the saliva, tear, and primary salivary gland epithelial cells (SGECs) of SS patients, as well as the salivary glands of the SS-prone non-obese diabetic (NOD/LtJ) mice. We also determined the functional roles of mt-dsRNAs in glandular acinar cell spheroids with dsRNA stress by measuring the SS glandular signature, such as ISGs and altered expression of AQP5/TJC proteins, and the inhibitory effect of Ach on mt-dsRNA induction with or without SS IgG. Endogenous dsRNA sensor for mt-dsRNA binding were validated and the regulators for mt-dsRNA induction were revealed by incubating the spheroids with inhibitors for JAK1 or mitochondrial RNA transcription.

## MATERIALS AND METHODS

### Saliva and tear samples

Tear samples were collected from the inferior tear meniscus using glass capillary tubes after applying 50 µl of normal saline in the inferior fornix and blinking for 5 sec. Schirmer test and ocular surface staining scoring were performed to assess tear production and the severity of ocular surface epithelial damage, respectively [24]. Saliva samples were collected as described previously [25]. Unstimulated and stimulated whole salivary flow rates were measured at the time of saliva sample collection. Detailed methods of saliva and tear sample collection and clinical characteristics of study participants are included in online supplementary material. The sample collection was approved by the Institutional Review Board (IRB, B-0506/021-004 and B-1806-472-301) of Seoul National University Bundang Hospital and all participants provided written informed consent.

### Preparation of human IgG

Human IgG was isolated from plasma samples of four patients with SS (SS-IgG) and four with rheumatoid arthritis (RA-IgG). The purified IgG fractions were pooled together (125 µg/patient) based on diagnosis. The patients with SS showed high titers of anti-M3R antibodies and those with RA were anti-M3R negative in our using our On-Cell-Western assay [26]. Further details are included in the online supplementary material.

### The salivary gland tissue collection from the NOD mouse

Female NOD/ShiLtJ mice (ages ranged from 8 weeks to 28 weeks, The Jackson Laboratory) were maintained under specific pathogen-free conditions. Mouse submandibular gland tissues were fixed in formalin and embedded in paraffin for immunohistochemistry and the tissue sections were dewaxed for RNA extraction. The RNA purification details, including the mouse primer sequences for RT-qPCR, available in Supplementary Appendix I (Table 5) and Appendix II (Materials and Methods). The study was approved by the Institutional Animal Care and Use Committee at the School of Medicine, The Catholic University of Korea (IACUC# CUMS-2019-0255-01).

### Cell culture and chemical treatment

SV40-immortalized NS-SV-AC cell was a kind gift from Dr. Masayuki Azuma (Department of Oral Medicine, University of Tokushima Graduate Faculty of Dentistry, Japan) [27]. Primary SGECs were obtained from primary SS patients, following the informed consent forms signed in accordance with the approved IRB protocol by the Institutional Review Board of Seoul St. Mary’s Hospital (IRB# KC13ONMI0646). The detailed protocols are included in Supplementary Appendix (Materials and Methods).

To induce dsRNA stress, the cells were transfected with 20 µg/ml of poly I:C (Sigma Aldrich) using Lipofectamine 3000 (Thermo Fisher Scientific) for 14 h following the manufacturer’s instruction, and Diethyl pyrocarbonate (DEPC)-treated water (RNase-free) was used for the control mock transfection. To test the effect of Ach or SS-IgG on mt-dsRNAs induction, 100 µM of Ach (Sigma Aldrich) was co-treated with poly I:C. SS-IgGs were also used with a control IgG from human serum (Sigma Aldrich). To examine the involvement of JAK1 in mt-dsRNA induction, a JAK1-inhibitor, 1 µg/mL of upadacitinib (MedKoo Biosciences), was treated 1 h prior to poly I:C transfection. To block the synthesis of mt-dsRNAs, cells were pretreated with 20 µM of 2-C′-methyladenosine (2-CM) (Santa Cruz Biotechnology) 24 h prior to poly I:C transfection.

### 3D spheroid culture on a pV4D4-coated plate

A 300 nm-thick pV4D4 film was deposited directly onto tissue culture polystyrene (TCPS) via the initiated chemical vapor deposition (iCVD) process and vaporized to be introduced to the iCVD chamber for monomer deposition and absorption. The detailed procedure is available in Supplementary Appendix II. Cells were seeded at the density of 1×10^5^ cells/ml on a pV4D4-coated plate and the spheroids formed on the plate were imaged at 24, 48, and 72 h by phase-contrast microscopy (Eclipse Ti-U; Nikon).

### RNA purification and RT-qPCR

RNA preparation, reverse transcription and RT-qPCR analysis were performed as previously described [11] and the details are included in Supplementary Appendix I (Tables S2-S4) for human primer sequences and Appendix II (Materials and Methods). To analyze the localization of mt-RNAs, cytosolic RNAs were isolated from cell lysates using the subcellular protein fractionation kit (Thermo Fisher Scientific) following the manufacturer’s instruction. TRIzol LS (Ambion) was added individually at a 3:1 volume ratio to the cytosolic fraction to extract RNAs.

### Protein expression analysis by western blotting

Western Blotting was performed as previously described [11], which is also included in Supplementary Appendix II. The primary antibodies, pPKR (Abcam, Ab81303), PKR (Cell Signaling Technology, D7F7), and β-tubulin (Cell Signaling Technology), for this study were used at a dilution of 1:1000, followed by secondary antibodies at a dilution of 1:5000.

### Cell viability measurement by acid phosphatase assay (APH)

APH assay for cell viability was performed as previously described [28] (Supplementary Appendix II).

### Immunocytochemistry/Immunohistochemistry and RNA single-molecule fluorescent *in situ* hybridization (RNA-FISH)

To analyze the expression of AQP5 and TJC protein expression, spheroids were transferred on a confocal dish (SPL). Following fixation and permeabilization, cells were incubated with primary antibodies: Occludin (Invitrogen), ZO-1 (Cell Signaling Technology), and AQP5 (Cell Signaling Technology) at a dilution of 1:100. Secondary antibodies were diluted at a 1:500 ratio, and fluorescent images were obtained using Zeiss LSM 780 confocal laser-scanning microscope with 63x (NA = 1.40) objective. The fluorescence intensity per area was quantified as integrated density divided by the area of each spheroid and then normalized to the control. See details in Supplementary Appendix II.

To visualize heavy and light strands of mt-Nd5 transcript from the NOD salivary gland tissue, five-micrometer sections of paraffin-embedded tissues were dewaxed using xylene, then dehydrated in ethanol for staining with RNAscope RNA fluorescent *in situ* hybridization (ACDBio) probes. Hybridization of the probe and signal amplification were performed with the manufacturer’s instructions.

### PKR formaldehyde-crosslinking immunoprecipitation (fCLIP)

To determine if mt-dsRNAs bind to cytosolic dsRNA sensor, fCLIP with anti-PKR antibody (Cell Signaling Technology) was performed. In brief, protein A beads (Thermo Fisher Scientific) were incubated with PKR antibody (Cell Signaling Technology). Harvested cells were fixed with 0.1% (w/v) paraformaldehyde (Sigma Aldrich) for 10 min. The crosslinked cells were lysed and the lysate was added to the PKR antibody-conjugated beads and incubated for 3 h at 4°C. The beads were washed and PKR-dsRNA complex was eluted from the beads. The eluate was treated with 2 mg/ml proteinase K (Sigma Aldrich) for overnight at 65°C and RNA was purified by acid-phenol:chloroform pH 4.5 (Thermo Fisher Scientific). The details for PKR fCLIP are included in Supplementary Appendix II.

### Statistical and functional analysis of mRNA-seq data

The read quality was checked with FastQC, and reads were aligned to the human genome (hg38) using Hisat2 (ver 2.1.0) [29]. The aligned reads were quantified with StringTie [30] using human gene annotation file version 32 from Gencode as the index. Differentially expressed genes (DEGs) were searched with DESeq2 [31], which was performed three times separately in each of DMSO and 2-CM-pretreated groups. Within each group, raw counts of two biological replicates with poly I:C transfection were compared against the raw counts of two replicates with the control group (i.e. DEPC transfection). Among genes that show log2 fold change with p-values less than 0.05, the values of DMSO (i.e. control) were subtracted from those of 2-CM-pretreated RNAs to evaluate the degree of rescue in the induction of gene expression. Top 200 of the genes that show rescue effects were assembled into a gene list for GO analysis by the ClueGo software (Cytoscape, v. 3.7.1) [32].

### Statistical Analysis

Continuous variables were analyzed using the one-tailed Student’s t-test or one-way ANOVA. Non-parametric test was performed using Mann-Whitney U Test. Receiver operating characteristic (ROC) analysis was performed to determine the area under the ROC curve of using mtRNA levels to distinguish SS from non-SS sicca. Bivariate correlations were analyzed using Pearson’s correlation coefficient. All data were biologically replicated at least three times. The error bars indicate the standard error of the mean. P values ≤ 0.05 were considered statistically significant. * denotes p-values ≤ 0.05, ** is p-values ≤ 0.01, and *** is p-values ≤ 0.001.

## RESULTS

### mt-dsRNA expression is elevated in SS patient saliva and tears, which was associated with secretory dysfunction and glandular inflammation

We first performed strand-specific reverse transcription to determine the level of heavy and light strand RNAs separately in the saliva and tear samples collected from 73 and 16 study subjects, respectively (Fig 1A). The heatmap showed higher expression salivary levels of most mtRNAs in SS patients, when compared to non-SS sicca patients or non-sicca controls. Salivary levels of *ND1L, ND4H, ND6H*, or *CYTB_H* were comparable among the three groups. Among 16 mtRNAs analyzed, saliva of SS patients showed significantly higher expression levels of 15 and 13 mtRNAs, when compared to non-SS sicca and non-sicca control patients (Fig 1B). It is notable that mtRNA expression patterns of non-sicca control and non-SS sicca group were indistinguishable (P>0.05). Additionally, we measured mt-dsRNA expression levels in 16 tear samples – 8 SS and 8 non-SS sicca subjects (Fig 1C) – and discovered increased levels of most mt-dsRNAs in tears of SS.

**Figure 1.**
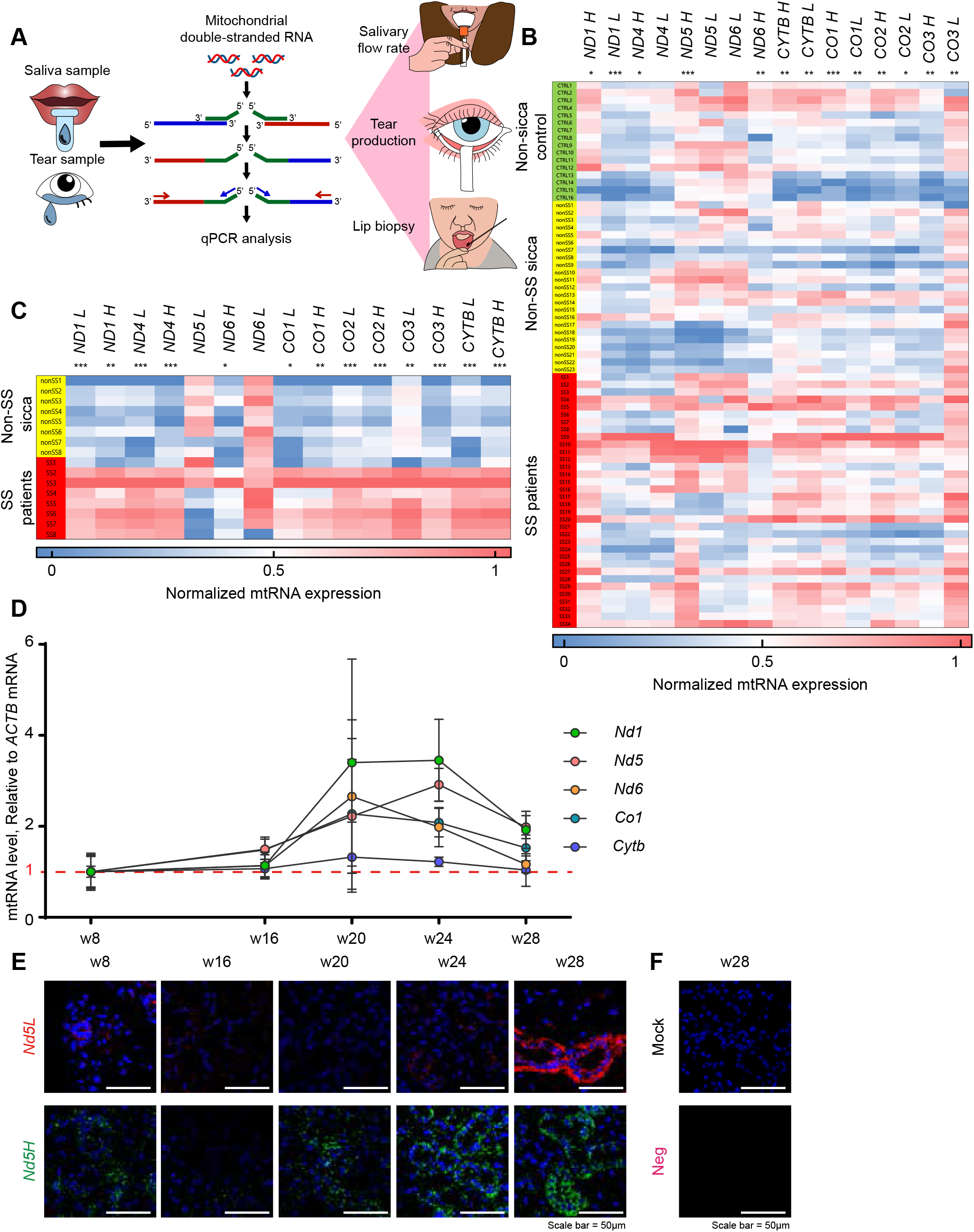
Elevated mt-dsRNA in saliva and tear of SS patients and the salivary glands of aged NOD mice. (A) RNA levels of mt-dsRNAs from the patient saliva and tear samples were measured, and the correlation between SS symptoms were analyzed. (B) Heatmap showing the relative RNA expression levels of both heavy (H) and light (L) strands in saliva among non-sicca control, non-SS sicca and SS patients. (C) Heat map showing the relative RNA expression levels in tears between non-SS sicca and SS patients. For clinical samples, p-values were calculated using ANOVA analysis, *p ≤ 0.05, **p ≤ 0.01, and ***p ≤ 0.001. (D) All RNA levels of five SS-associated mt-dsRNAs in the salivary gland of NOD mice. Cq values are relative to *GAPDH* mRNA, then normalized to non-sicca and non-SS sicca controls. Results are presented as the mean ± SEM of three independent samples. (E) Representative fluorescence images of NOD mouse salivary gland tissues with mt-Nd5 light (*Nd5L*) and heavy strand (*Nd5H*) transcripts. (F) The negative control probe yielded no signal. The scale bars represent 50 µm.

When the five most abundant mitochondrial transcripts in tear and saliva samples were ranked (Fig S1A and B, respectively), *CYTB_H ND1L*, and *CO1L* expressions were highly upregulated in SS compared to non-SS sicca patients in both samples. Interestingly, tear *CYTB_L* (Fig S1C-E) and salivary *ND5H* (Fig S1F-I) levels were positively correlated with secretory dysfunction and the degree of tissue damage in SS.

### mt-dsRNA expression is upregulated in the salivary glands of the diseased SS-prone mouse

Aged NOD mice developing SS-like autoimmune exocrinopathy is one of the most extensively studied animal models in deciphering SS pathogenesis [33]. Since there is a significant decrease in the salivary flow rate of NOD mice after 17 weeks with a full-blown disease phenotype [34], we focused our analysis on 16 to 28-week-old NOD mice compared to 8-week-old mice as controls. Total RNA was extracted from the tissue sections of the NOD salivary glands was subject to RT-qPCR for *Nd1, Nd5, Nd6, Co1*, and *Cytb*, which showed the most significant enrichment in the saliva samples of SS patients. Compared to that of 8-week-old NOD mice, expression of all mt-dsRNAs except *Cytb* were higher in NOD older than 20 weeks (Fig 1D). However, due to high variabilities in the values, the ANOVA analysis did not yield any significance despite a clear increase in the mean expression of the RNAs of interest. Alternatively, we analyzed mt-dsRNA patterns by using RNA-FISH. The expression of both *Nd5H* and *Nd5L* RNAs were clearly elevated in the acinar regions of diseased NOD submandibular salivary gland tissues (Fig 1E), compared to the level detected in younger mice, implying the increased expression of mt-dsRNAs. Note, the negative control probe yielded no signal (Fig 1F).

### dsRNA induces mt-dsRNAs in the salivary gland acinar cell line and primary SGECs

At the next step, we investigated whether stimulating the antiviral signaling and I-IFN pathway in SGECs can elicit mt-dsRNA induction. In order to induce SS-like condition, we transfected SV-40 transformed salivary gland-squamous acinar cells (NS-SV-AC) with poly I:C, a synthetic analog of viral dsRNAs. Repeated administration of poly I:C was reported to elicit IL-7 production and early inflammatory responses in the salivary gland and accelerate the development of SS-like sialadenitis [35, 36]. Moreover, a study indicates that poly I:C leads to other key characteristics of SS, such as leukocytic infiltration, antinuclear antibody production, and impaired tear secretion in mice [37]. As shown in Fig S2A-C, poly I:C stimulation resulted in robust induction of ISGs, increased PKR phosphorylation, and consequent decrease in cell viability.

We then measured the mt-dsRNA induction upon poly I:C stimulation. Consistent with results in patient salivary and tear samples, we found that dsRNA stress led to increased mt-dsRNA expression, both heavy and light strand RNAs (Fig S2D and E). Direct visualization of *ND5* mt-dsRNA via RNA-FISH further confirmed that the expression of both heavy and light strand RNAs were increased (Fig S2F). The isolated RNAs from the free cytosolic compartment were analyzed, revealing increased mt-dsRNAs in the cytoplasm (Fig S2G). We also employed fCLIP to examine the interaction between the mt-dsRNAs and PKR. Our data confirmed the enhancement of PKR-mtRNA interaction upon poly I:C stimulation (Fig S2H). Considering that both poly I:C and mt-dsRNAs can activate PKR, our results suggest that increased cellular mt-dsRNAs may amplify the effect of dsRNA-mediated antiviral signaling.

To further demonstrate that mt-dsRNA induction by poly I:C is not limited to the stable cell line, we examined the mt-dsRNA expression in the SS primary SGECs. We found that SGECs of SS patients tended to exhibit higher mt-dsRNA expression levels compared to those from non-SS sicca patients (Fig S3A). Moreover, poly I:C stimulated resulted in mt-dsRNA induction in primary cells of SS and non-SS sicca subjects (Fig S3B and C), similar to the induction in the NS-SV-AC cells.

### dsRNA stress downregulates AQP5 and TJC proteins in 3D microenvironment, concurrently with mt-dsRNA induction

Altered expression/distribution of TJC and AQP5 proteins could lead to secretory dysfunction of salivary and lacrimal glands, both qualitatively and quantitatively [21-23, 38, 39]. Because pV4D4-coated surface-based spheroid culture (Fig 2A) made more homogeneous in terms of both shape and size than 2D cultures or Ultra-Low attachment plate (ULA), the 3D spheroid salivary gland cells were able to clearly analyze the protein expression in immunofluorescence imaging analysis (Fig S4A-S4C).

**Figure 2.**
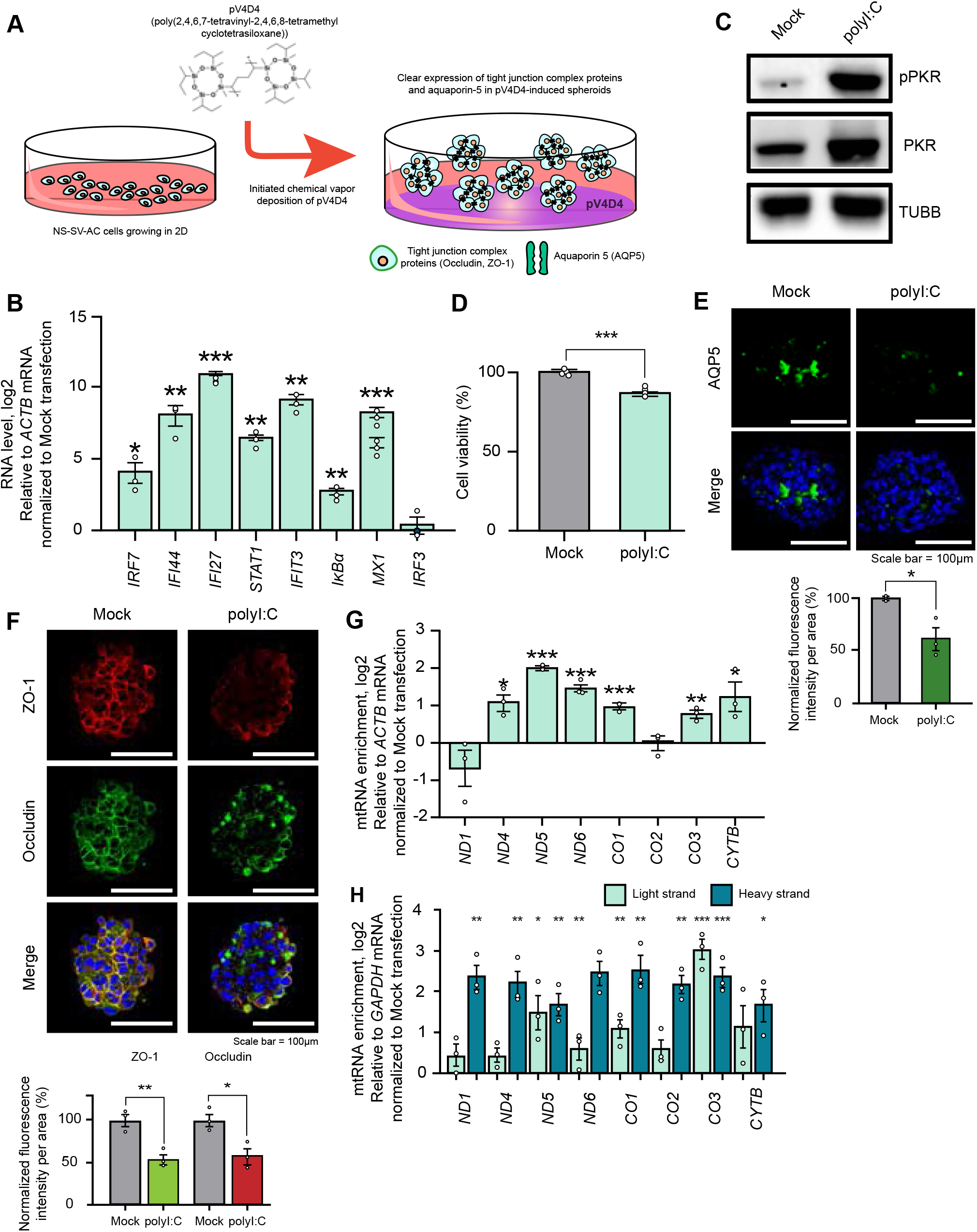
mt-dsRNA induction and the SS glandular features by dsRNA stimulation in salivary acinar cell spheroids. (A) A schematic representation of 3D cell culture by depositing V4D4 monomers via iCVD process. (B) Relative mRNA expression of indicated ISGs in poly I:C transfected NS-SV-AC spheroids. The normalized log_2_ fold enrichment of three independent experiments. (C) Western blot analysis of PKR phosphorylation upon poly I:C transfection in 3D cell culture system. (D) APH assay showing cell viability upon poly I:C transfection (n = 4). (E) Representative fluorescence images showing reduced expression of AQP5 and (F) TJC proteins upon poly I:C transfection. The fluorescence intensities were quantified as the mean of three independent experiments and normalized to the mock transfection control group. The scale bars represent 100 µm. (G) Relative RNA level of total mtRNAs and (H) stand-specific mtRNAs in poly I:C transfected NS-SV-AC spheroids. The normalized log_2_ fold enrichment of three independent experiments. Unless mentioned otherwise, Cq values are relative to *ACTB* mRNA, then normalized to the mock transfected control (B, G, and H). All error bars represent SEM, and p-values were calculated using one-tailed Student’s t-tests, *p ≤ 0.05, **p ≤ 0.01, and ***p ≤ 0.001.

Upon poly I:C stimulation to 3D spheroid salivary gland cells, consistent with our previous results in 2D culture, ISGs and PKR phosphorylation were increased, and a decrease in cell viability was evident (Fig 2B-D). Poly I:C stimulation also led to a dramatic decrease in AQP5 (Fig 2E), and reduction in ZO-1 and Occludin (Fig 2F). Most importantly, we confirmed that poly I:C stimulation led to induction of mt-dsRNAs (Fig 2G-H).

### Effects of Ach are mediated via mt-dsRNAs

Considering that Ach, the endogenous ligand for M3R, can block cytosolic export of mtDNAs [40], we investigated whether Ach affects the expression of mt-dsRNAs in salivary gland cells. We found that, in the presence of Ach, ISGs induction was attenuated, and this phenomenon was accompanied by decreased induction of mt-dsRNAs (all p<0.05; Fig 3A and B). To investigate the effect of mt-dsRNAs on Ach-induced suppression of ISGs, we depleted mt-dsRNAs using 2-CM, an inhibitor of POLRMT. Pretreatment of 2-CM led to a dramatic decrease in the expression of all mt-dsRNAs examined (Fig 3C). Interestingly, we found that Ach-induced ISGs suppression was significantly diminished in mt-dsRNA-depleted cells (all p<0.05; Fig 3D), thereby suggesting that the suppression of ISGs by Ach was partly mediated by mt-dsRNAs.

**Figure 3.**
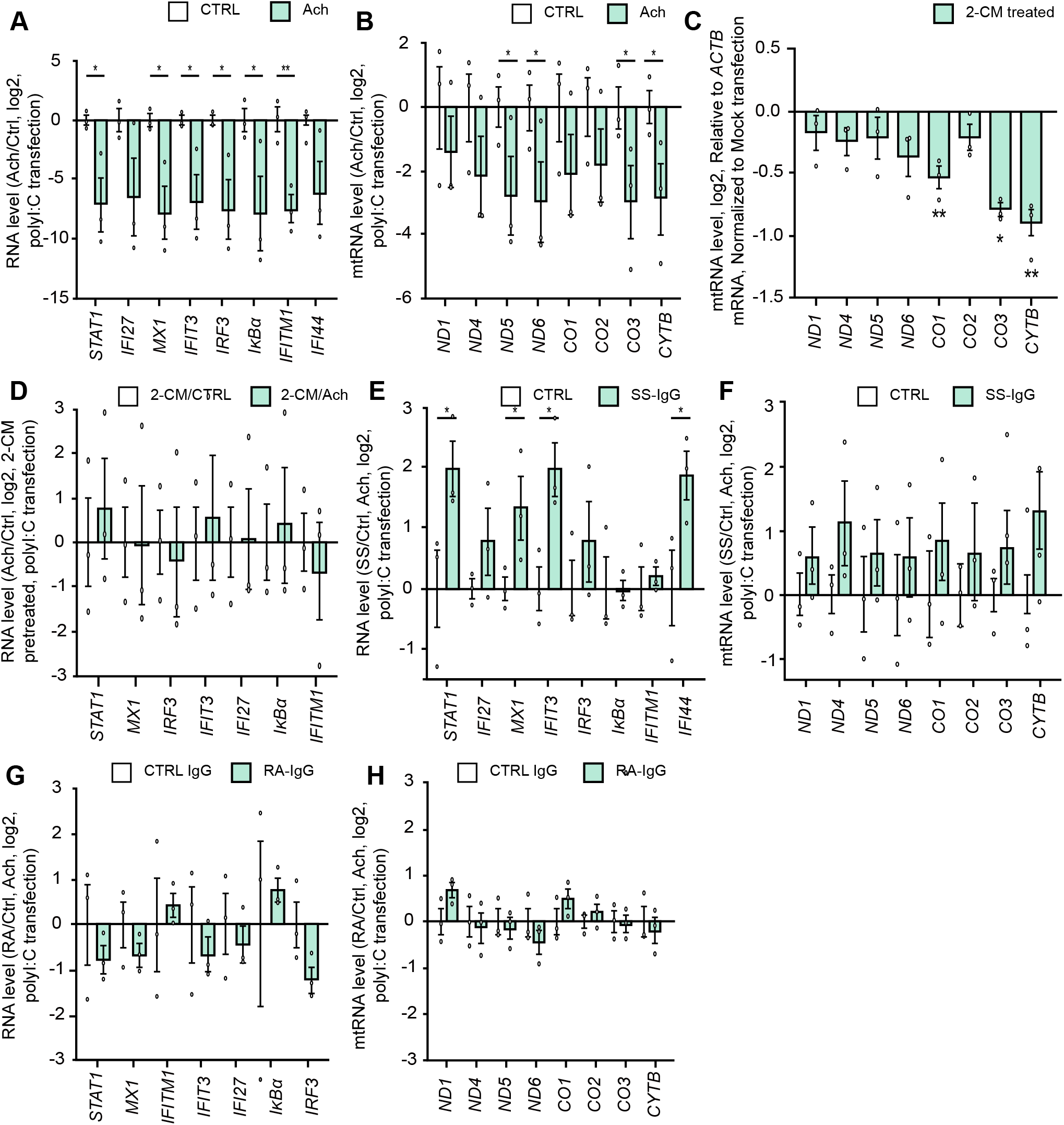
M3R ligand Ach preventing mt-dsRNA induction to yield the protective effects against SS-like salivary gland cell phenotypes. (A) The ratio of ISGs and (B) mt-dsRNAs expression upon poly I:C transfection in the presence or the absence of 100 µM Ach. (C) Reduced expression of mt-dsRNAs upon pretreatment of 20 µM 2-CM in NS-SV-AC spheroids. (D) The ratio of ISGs induction upon poly I:C transfection in Ach-treated samples after 2-CM pretreatment. (E) The ratio of ISGs and F) mt-dsRNAs expression upon poly I:C stimulation in the presence of 100 µM Ach and 500 µg of control IgG or SS-IgG. (G) The ratio of ISGs and (H) mt-dsRNAs expression upon poly I:C stimulation in the presence of Ach with either control IgG or RA-IgG. All Cq values are relative to *ACTB* mRNA. The values of poly I:C-transfected samples were normalized to the mock-transfected samples of the same DMSO- or the Ach-treated group. Unless mentioned otherwise, the graph represents three independent experiments with error bars denoting SEM. All statistical significances were calculated using one-tailed Student’s t-tests, *p ≤ 0.05, **p ≤ 0.01, and ***p ≤ 0.001.

We next analyzed whether SS-IgG containing anti-M3R autoantibodies could counteract the Ach-induced ISG suppression. When spheroid salivary gland cells were treated with either control IgG or SS-IgG, SS-IgG resulted in significantly increased ISGs induction and mt-dsRNA expression, thereby lessening the effect of Ach (all p<0.05; Fig 3E-3F). However, RA-IgG that do not contain anti-M3R autoantibodies did not affect the Ach-induced suppression of ISG and mt-dsRNA expression (Fig 3G-H).

### mt-dsRNA and ISG induction by dsRNA stress in the spheroids depends on JAK1

Considering the concomitant induction of mt-dsRNA with ISG expression and the replication of SS glandular signatures presented in this study, we hypothesize that mt-dsRNA induction is contingent upon the Janus kinase (JAK)-signal transducer and activator of transcription (STAT) pathway as ISGs are known to be unregulated in response to I- and II-IFNs via JAK1 [41, 42]. NS-SV-AC spheroids were pretreated with upadacitinib, a JAK1 inhibitor, before poly I:C stimulation, which resulted in significant reduction of ISGs and mt-dsRNAs (all p<0.05; Fig 4A and 4B). Moreover, 10 μg/mL of upadacitinib treatment resulted in decreased pPKR (Fig 4C) and increased cell viability (Fig 4D). Lastly, we found that upadacitinib prevented the poly I:C-induced downregulation of AQP5 and TJC proteins (Fig 4E-F).

**Figure 4.**
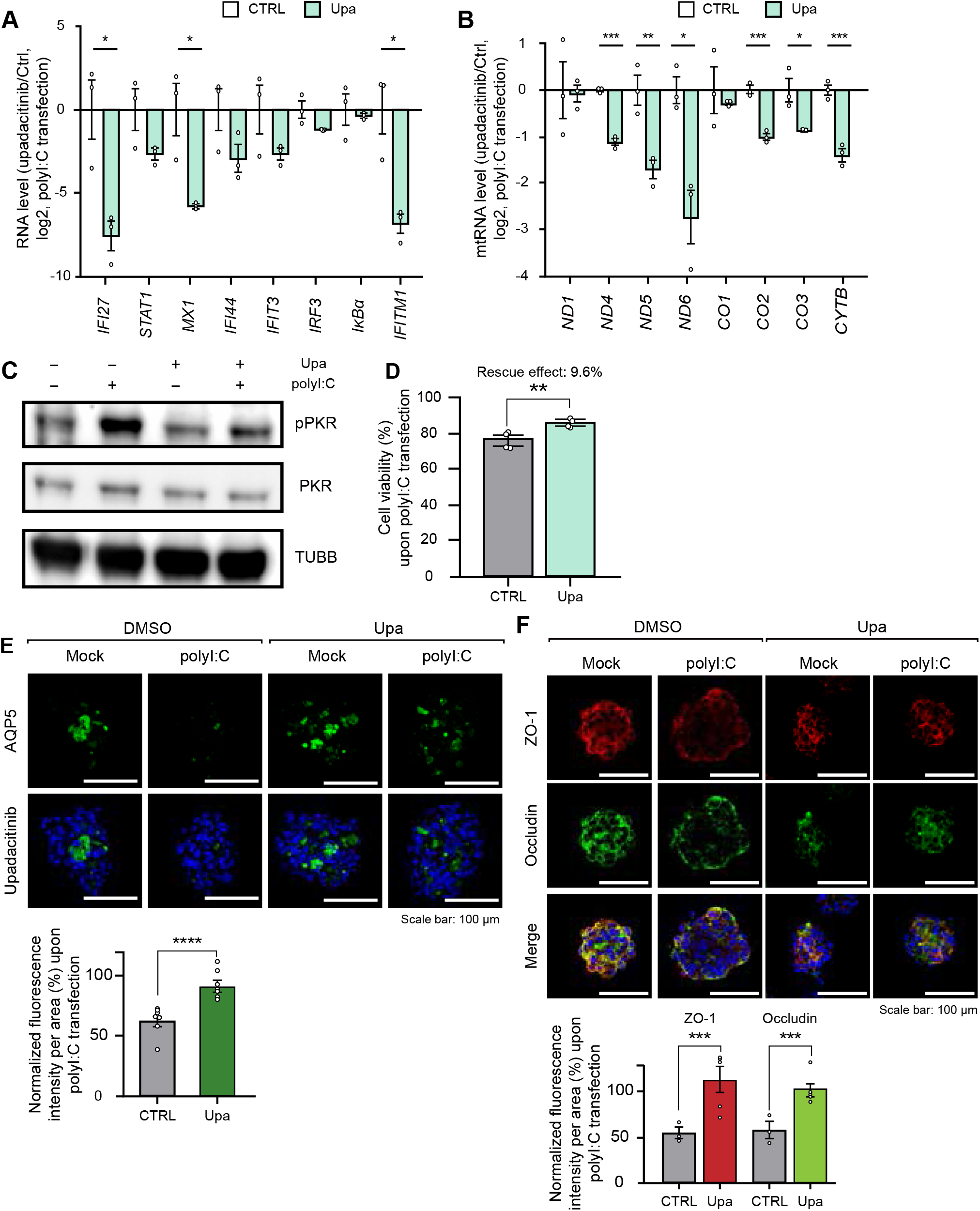
mt-dsRNA induction mediated by the JAK1 signaling pathway. (A) The ratio of ISGs and (B) mt-dsRNAs expression upon poly I:C transfection with or without upadacitinib treatment. All Cq values are relative to *ACTB* mRNA. The values of poly I:C-transfected samples were normalized to the mock-transfected samples of the same DMSO- or the upadacitinib-treated group. (C) Western blot analysis of rescue effect in PKR phosphorylation and (D) cell viability measured by APH assay when upadacitinib is treated prior to poly I:C transfection. (E) Representative fluorescence images showing the rescue effect in AQP5 and (F) TJC protein expression upon upadacitinib pretreatment prior to poly I:C transfection. The fluorescence intensities were quantified as the mean of three independent experiments and normalized to the mock transfection control group. The scale bars represent 100 µm. Unless mentioned otherwise, three independent experiments were carried out, and error bars denote SEM. All statistical significances were calculated using one-tailed Student’s t-tests, *p ≤ 0.05, **p ≤ 0.01, and ***p ≤ 0.001.

### Downregulating mt-dsRNA synthesis attenuates the I-IFN signature

In order to confirm the impact of mt-RNAs on IFN-mediated immune regulation and the SS glandular signature, we analyzed poly I:C treated cells with 2-CM via mRNA-seq. We found that 2-CM pretreatment significantly decreased the expression of genes related to the I-IFN signaling pathway (Fig 5A). Half of the number of I-IFN signaling genes showed reduced levels, and such results were further confirmed using RT-qPCR (Fig S5A). While poly I:C transfection still induced ISGs, the degree of induction was significantly reduced when mt-dsRNAs were downregulated, suggesting that the increased expression of mt-dsRNAs and their cytosolic export may aggravate the glandular cellular response triggered by poly I:C stimulation. The GO analysis revealed that genes related to oxidative phosphorylation (4.55%) and negative regulation of viral genome replication (18.18%) were partly rescued upon 2-CM pretreatment (Fig S5B). Moreover, 2-CM resulted in decreased induction of genes involved in lymphocyte chemotaxis (3.03%).

**Figure 5.**
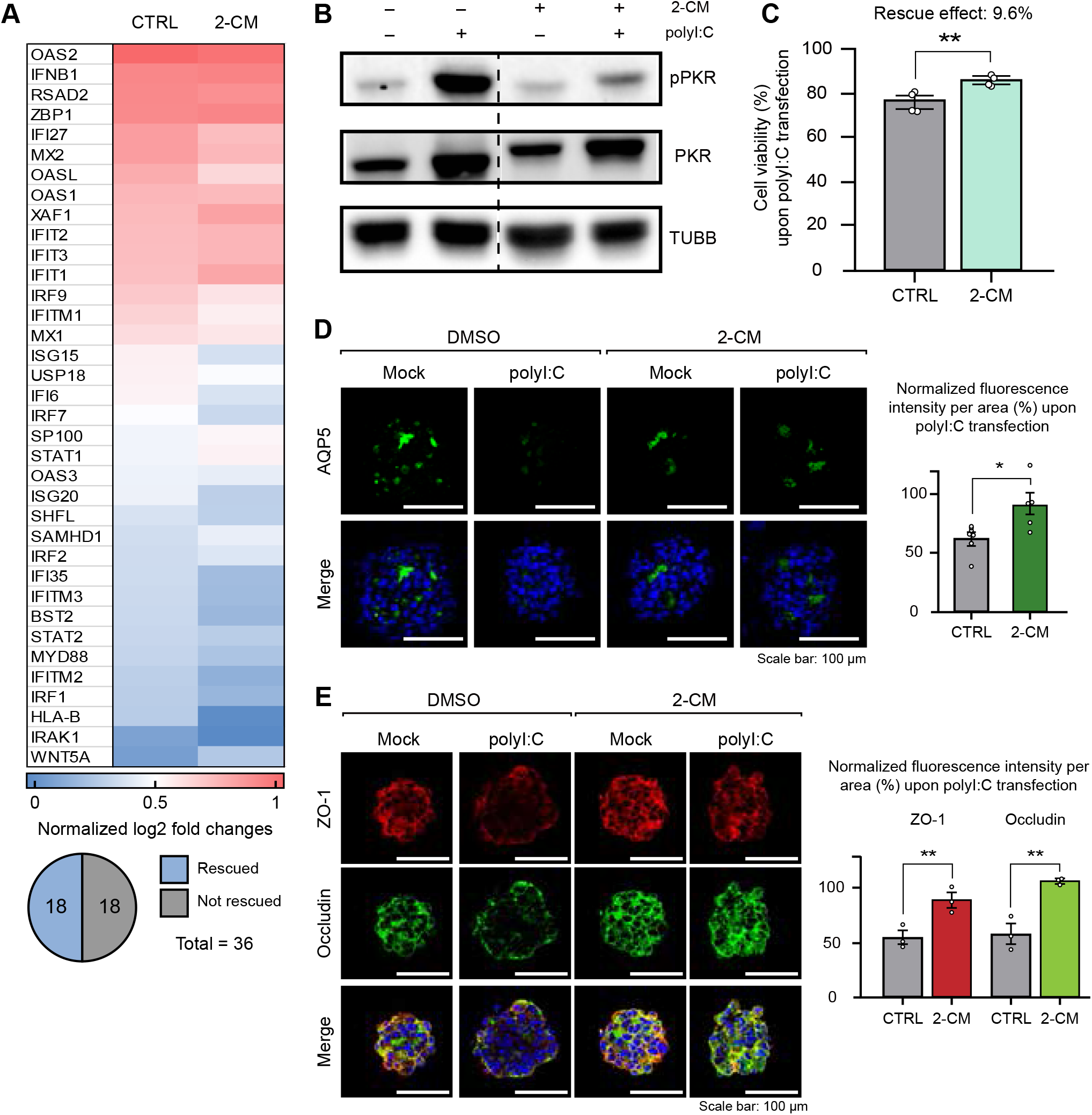
mt-dsRNA downregulation attenuating the cellular response to poly I:C stimulation. (A) Heatmap of DEG analysis for type I IFN signaling pathway genes. The two columns represent log_2_ fold changes with or without 2-CM pretreatment 24 h prior to poly I:C stimulation. (B) Western blot analysis of rescue effect in PKR phosphorylation and (C) cell viability measured by APH assay upon downregulation of mt-dsRNAs before poly I:C transfection. (D) Representative fluorescence images of AQP5 and (E) TJC proteins–ZO-1 and Occludin-upon poly I:C stimulation with or without 2-CM pretreatment. The fluorescence intensities in were quantified as the mean of three independent experiments and normalized to the mock transfection control group. The scale bar represents 100 µm. All statistical significances were calculated using one-tailed Student’s t-tests, *p ≤ 0.05, **p ≤ 0.01, and ***p ≤ 0.001.

Downregulation of mt-dsRNAs also reduced phosphorylation of PKR (Fig 5B), and along with decreased ISGs expression, this consequently rescued cell viability by approximately 10% (Fig 5C). Furthermore, 2-CM pretreatment counteracted the negative effect of poly I:C on the expression of AQP5, ZO-1, and Occludin (Fig 5D-5E).

## DISCUSSION

Our study demonstrated for the first time that 1) mt-dsRNAs were characteristically elevated in saliva and tear samples of SS patients and were associated with exocrine dysfunction and inflammation in the lip biopsy specimen; 2) mt-dsRNAs were upregulated in MSG-derived SGEC from SS patients; 3) mt-dsRNAs were increased in salivary glands of SS mouse model; 4) Ach-induced suppression of poly I:C-mediated changes was mediated partly by mt-dsRNAs, and SS-IgG negated such protective action of Ach; 5) mt-dsRNA induction was under the control of the JAK1 pathway; 6) downregulation of mt-dsRNA ameliorated the SS-related cellular response in salivary gland cells (Fig 6). These results suggest that mt-dsRNAs could be a novel mediator in the pathophysiology of exocrinopathy underlying SS.

**Figure 6.**
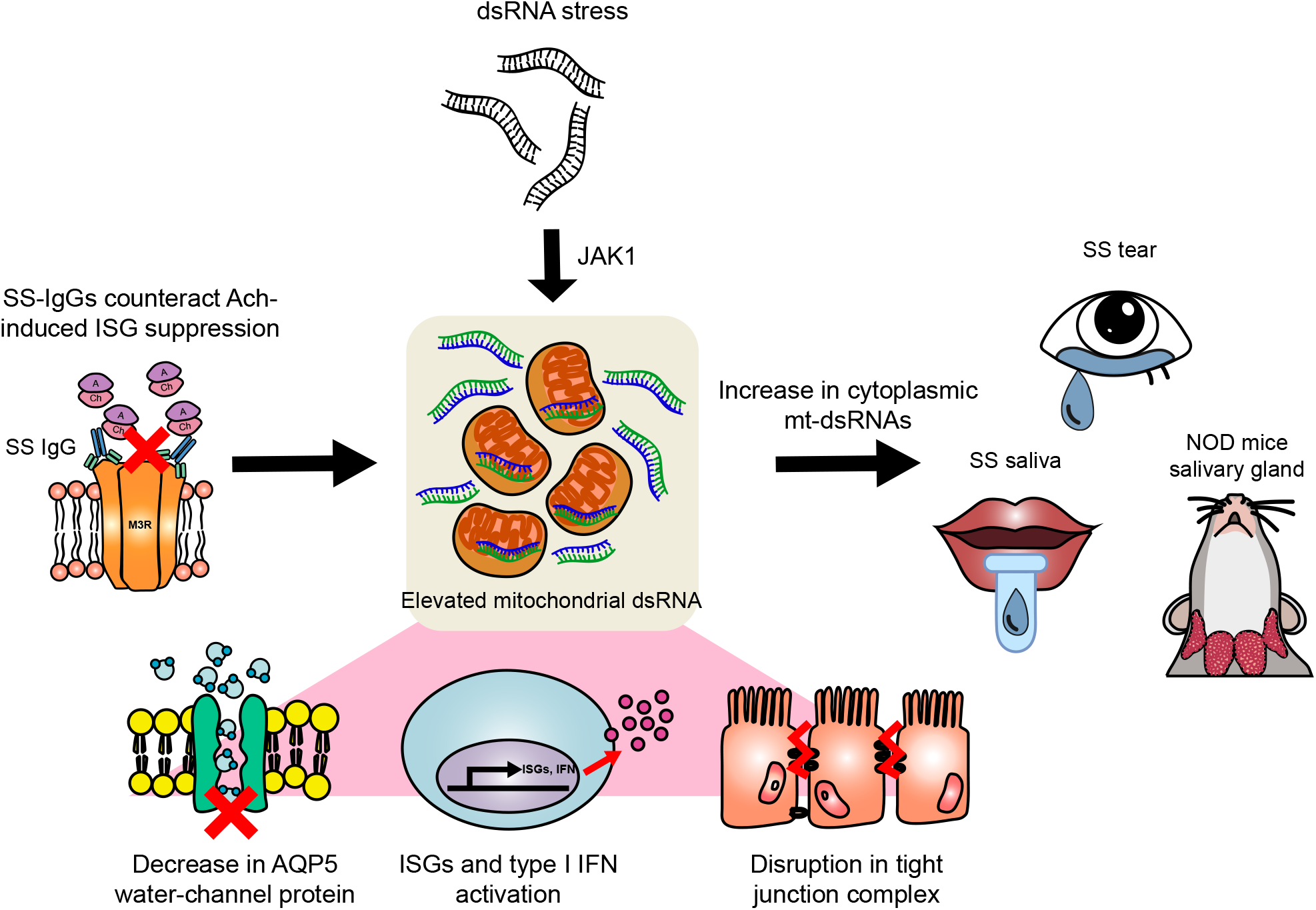
mt-dsRNAs as the molecular amplifier of the glandular autoimmune phenotypes in SS. dsRNA stress leads to elevation of mt-dsRNAs via the JAK1/STAT pathway. SS-IgGs also results in mt-dsRNAs induction, thereby lessening the effect of Ach. Accumulation of mt-dsRNAs contributes to exacerbation of SS-related phenotypes in the salivary gland acinar cells. Elevated levels of mt-dsRNAs are detected in both human samples and the SS-prone mouse model. Countering the enrichment of mt-dsRNAs may alleviate the pathological changes reported in the glands of SS.

Nucleic acid sensing by PRRs occurs in non-immune cells as well as professional immune cells to induce anti-viral responses including secretion of various alarmins and cytokines and cell death. Chronic inflammation can result from aberrant activation of IFNs, and the persistent activation of IFN signaling is considered to the pathogenesis of SS, although both IFN pathway activation and genetic association with IFN signaling are shared by other systemic autoimmune diseases. Our study provides a potential link between cytosolic mitochondria-derived dsRNAs and the activation of PRRs, which subsequently leads to upregulation of I-IFNs. Mitochondrial dysfunction and accumulation of damaged mitochondria are frequently reported in SS patients [43, 44]. Additionally, it has been reported that poly I:C induced reactive oxygen species (ROS) stress and disruption of mitochondrial potential in SGECs [45, 46]. We found that poly I:C stimulation resulted in the cytosolic release of mt-dsRNAs, and subsequent PKR phosphorylation and ISG induction. Notably, downregulation of mt-dsRNAs via 2-CM resulted in the attenuation of IFN signature. Therefore, mt-dsRNAs could be another endogenous ligand of cytosolic RNA-sensing pathway, apart from retroelements and non-coding RNAs [7].

In the present study, mt-dsRNAs were detected in saliva and tear samples by RT-qPCR. mt-dsRNAs can be exported outside the cells and work as extracellular signaling molecules [14, 15]. Interestingly, we observed that extra-cellular mt-dsRNAs levels were elevated in the body fluids of SS patients and their levels were associated with exocrine dysfunction or damage and lymphocyte infiltration. Although the origin remains to be elucidated, salivary or tear mt-dsRNAs can be a potential diagnostic biomarker with pathophysiological background in SS.

Ben *et al*. reported that Ach prevents cytoplasmic mtDNA release through mitochondrial α7 nicotinic Ach receptor (α7nAChR), leading to the inhibition of NLRP3 inflammasome [47]. In this study, Ach also suppressed the effects of poly I:C via the modulation of cytosolic mt-dsRNAs levels in salivary gland cells. It was reported that the number of the nerve terminals is reduced in the MSG of SS patients and parasympathetic nerve fibers are decreased in areas of focal infiltrates in NOD mice [47, 48]. SS-IgG contains antibodies reacting with M3R and inhibits the function of salivary acinar cells [49, 50]. SS-IgG were prepared from SS patients with anti-M3R, which were determined by our cell-based method [24], and it negated the protection conferred by Ach in salivary gland cells in this study. Additionally, the presence of anti-M3R in SS patients could explain the decrease in response to M3R agonists in those with hypergammaglobulinemia [51]. Therefore, salivary glands in SS patients have topographic or functional interference of Ach signaling that provides a supportive environment. In spite that a peak effect of pilocarpine on salivary flow starts within 1 h in healthy volunteers, its therapeutic benefit may be delayed until after 12 weeks of treatment [52]. In the salivary glands of a SS murine model based on NFS/sld mice, M3R expression was reported to be elevated after 2-week administration of pilocarpine [57]. Considering together with our findings showing the protective action of Ach, continuous M3R stimulus could restore secretory function, at least partially, in patients with SS-related hyposalivation. The interrelationship of mt-dsRNAs with M3R should be further investigated to elucidate how the aberrant autoimmune response leads to mt-dsRNA-mediated SS-related changes of salivary gland cells.

Another novel finding of this study is that mt-dsRNA induction is under the control of JAK1. Poly I:C-induced IFNs activate heterodimers of JAK1 via IFN receptors; JAK1/TYK2 for I-IFNs and JAK1/JAK2 for II-IFN. In the present study, upadacitinib, a JAK1 inhibitor, decreased poly I:C-induced mt-dsRNA induction and downregulation of AQP5 and TJC proteins in salivary gland cells. Previously, JAK inhibitors reported to suppress IFNγ-induced CXCL10, ROS-induced STAT3 activation, and IFN-induced BAFF secretion in salivary gland cells [41, 53, 54]. Therefore, our findings could serve as another basis for clinical trials of JAK inhibitors in SS.

The limitations of the current study include unknown cellular origin of salivary and tear mt-dsRNAs although we presume to be the salivary and lacrimal glands. However, involvement of apoptotic or desiccating buccal, gingival, corneal, conjunctival, and even immune cells through inflamed micro-vessels cannot be ruled out. Because we focused on cell-intrinsic mt-dsRNA sensing via PKR, it is unclear if mt-dsRNAs in saliva or tear can generate similar impact on the cells as what is shown in our *in vitro* system. Recently Bonekamp et al. developed first-in-class inhibitors targeting the human mitochondrial RNA polymerase, LDC195943 and LDC203974 [55]. In future, a novel class of therapeutic agents could be investigated. Regardless, our novel discovery of upregulated mt-dsRNAs offers a potential to develop a non-invasive diagnostic tool, which can be clinically utilized in the form of liquid biopsy at the point-of care.

In summary, compared to extensive studies available on the roles of mt-dsDNA in autoimmune disorders, the roles of mt-dsRNAs have not been clearly elucidated. This current study underscores the critical roles of mt-dsRNAs in potentiating the SS glandular signatures in glandular spheroids under dsRNA stress by facilitating the positive feedback loop of the PKR, JAK1, and ISGs. Minimizing mt-dsRNA production by eliminating excessive mt-dsRNAs or accelerating mt-dsRNA decay can be a potential therapeutic target in SS, especially, in the context of glandular preservation prior to irreversible damage.

## Online supplemental material

Online supplemental material is available.

## Data and materials availability

All data needed to evaluate the conclusions in the paper are present in the paper and/or the Supplementary Materials. Additional data related to this paper may be requested from the first and/or corresponding author/s. Sequencing data that support the findings of this study are available in the GEO database under the accession number GSE168148. All other relevant datasets used and/or analyzed during the current study are available within the article and its Supplementary Materials files or from the corresponding authors on reasonable request. The computational pipeline used in this study is open-sourced and available at: https://daehwankimlab.github.io/hisat2/, https://ccb.jhu.edu/software/stringtie/, and http://www.bioconductor.org/packages/release/bioc/html/DESeq2.html.

## Acknowledgments

We thank all members of the Lee, Im, and Kim laboratory for helpful discussion and comments on the paper.

## Author contributions

J. Yoon, M. Lee, YJ. Lee, SG. Lim, and Y. Kim designed the study and analysis. Experiments were performed by J. Yoon, and M. Lee performed imaging and cell viability analysis of 3D spheroids. YJ. Lee provided the saliva samples. A.A. Ali analyzed the saliva samples and performed the initial characterization of 3D spheroids. YR. Oh and YS. Choi collected and prepared patient samples and developed a hAQP5-expressing NS-SV-AC cell line.

N. Lee processed the mRNA-seq raw data file for DEG analysis. SG. Jang and S. Kwok provided the NOD mouse tissue samples and primary cells of SS patients. J. Yoon and S. Kim analyzed the salivary gland tissues of NOD mice. JY. Hyon and S. Cha designed the project with YJ. Lee. The study was supervised by Y. Kim, YJ. Lee, and SG. Im. J. Yoon, YJ Lee, S. Cha, and Y. Kim wrote the manuscript with contributions from S. Kim. All authors subsequently reviewed and edited the manuscript.

