## Supplemental material for "Mitochondrial double-stranded RNAs as a pivotal mediator in the pathogenesis of Sjögren’s syndrome"

Additional Supporting Information may be found in the online version of this article at the publisher's website.

**SUPPLEMENTARY APPENDIX**

**Supplementary Table 1.** Clinical characteristics of female subjects providing tear or saliva samples.

**Supplementary Table 2.** Primer sequences for strand-specific reverse transcription.

**Supplementary Table 3.** Primer sequences for qRT-PCR.

**Supplementary Table 4.** Primer sequences for strand-specific qRT-PCR.

**Supplementary Table 5.** Primer sequences for qRT-PCR in mouse samples

**Supplementary Materials and Methods**

**Supplementary Figure 1.** Association of the five most significant mitochondrial RNA expression levels in body fluid with exocrine dysfunction.

**Supplementary Figure 2.** dsRNA stimulation leads to increased mt-dsRNA expression.

**Supplementary Figure 3.** Poly I:C stimulation leads to mt-dsRNA induction in primary SGECs isolated from SS patients.

**Supplementary Figure 4.** The 3D culture system enhances the expression of TJC and AQP5 proteins.

**Supplementary Figure 5.** mRNA-seq data validation using qRT-PCR and GO analysis of 2-CM-pretreated samples.

**Supplementary Table 1.** *Clinical characteristics of female subjects providing tear or saliva samples.*

| | Subjects donated tear samples | | Subjects donated saliva samples | | |
|---|---|---|---|---|---|
| | Non-SS sicca (n=8) | SS (n=8) | Non-sicca controls (n=16) | Non-SS sicca (n=23) | SS (n=34) |
| Age (years) | 66.4±2.7 | 60.9±3.3 | 42.5±1.9 | 47.1±2.1 | 46.0±2.0 |
| Dry eye symptoms* | 8/8 (100%) | 8/8 (100%) | | 21 (91.3%) | 29 (85.3%) |
| Dry mouth symptoms* | NA | 8/8 (100%) | NA | 16 (69.6%) | 25 (73.5%) |
| Lacrimal dysfunction* | NA | 8/8 (100%) | NA | 5 (21.7%) | 23 (67.6%) |
| Salivary dysfunction* | NA | 8/8 (100%) | NA | 8 (34.8%) | 29 (85.3%) |
| Anti-Ro/SSA positivity | NA | 8/8 (100%) | NA | 2 (8.7%) | 31 (91.2%) |
| Focus score ≥ 1 on minor salivary gland biopsy | NA | 2/4 (50%) | NA | 0/14 (0%) | 21/29 (72.4%) |
| Schirmer test (mm)§ | 16.8±3.7 | 7.5±5.2 | NA | NA | NA |
| Ocular staining score§ | 0.6±0.3 | 7.4±1.0 | NA | NA | NA |
| Whole salivary flow rate (mL/min)§ | | | | | |
| Unstimulated | NA | NA | NA | 0.228±0.042 | 0.124±0.016 |
| Stimulated | NA | NA | NA | 1.147±0.101 | 0.623±0.076 |
| Extra-glandular manifestations⁋ | - | 2 (25.0%) | - | - | 17 (50.0%) |

*, defined according to the American-European Consensus Group classification criteria for Sjögren's syndrome; §, determined at the collection site and time; ⁋, included arthritis (*n* = 11), hematologic abnormalities (*n* = 8), Raynaud's phenomenon (*n* = 8), peripheral neuropathy (*n* = 2), glomerulonephritis (*n* = 1), cutaneous lupus erythematosus (*n* = 1), and lymphocytic interstitial pneumonia (*n* = 1); SS = Sjögren's syndrome; NA = not available. Data were expressed as mean ± s.e.m. or number (percentage).

**Supplementary Table 2.** *Primer sequences for strand-specific reverse transcription.*

| Primer | Sequences (5′-3′) |
|---|---|
| ND1 Heavy | CGCAAATGGGCGGTAGGCGTGGTTGTGATAAGGGTGGAGAGG |
| ND1 Light | CGCAAATGGGCGGTAGGCGTGTCAAACTCAAACTACGCCCTG |
| ND4 Heavy | CGCAAATGGGCGGTAGGCGTGTGTTTGTCGTAGGCAGATGG |
| ND4 Light | CGCAAATGGGCGGTAGGCGTGCCTCACACTCATTCTCAACCC |
| ND 5 Heavy | CGCAAATGGGCGGTAGGCGTGTTTGGGTTGAGGTGATGATG |
| ND5 Light | CGCAAATGGGCGGTAGGCGTGCATTGTCGCATCCACCTTTA |
| ND6 Heavy | CGCAAATGGGCGGTAGGCGTGGGTTGAGGTCTTGGTGAGTG |
| ND6 Light | CGCAAATGGGCGGTAGGCGTGCCCATAATCATACAAAGCCCC |
| CO1 Heavy | CGCAAATGGGCGGTAGGCGTGTTGAGGTTGCGGTCTGTTAG |
| CO1 Light | CGCAAATGGGCGGTAGGCGTGGCCATAACCCAATACCAAACG |
| CO2 Heavy | CGCAAATGGGCGGTAGGCGTGGTAAAGGATGCGTAGGGATGG |
| CO2 Light | CGCAAATGGGCGGTAGGCGTGCTAGTCCTGTATGCCCTTTTCC |
| CO3 Heavy | CGCAAATGGGCGGTAGGCGTGCTCCTGATGCGAGTAATACGG |
| CO3 Light | CGCAAATGGGCGGTAGGCGTGCCTTTTACCACTCCAGCCTAG |
| CYTB Heavy | CGCAAATGGGCGGTAGGCGTGGGATAGTAATAGGGCAAGGACG |
| CYTB Light | CGCAAATGGGCGGTAGGCGTGCAATTATACCCTAGCCAACCCC |
| GAPDH | CGCAAATGGGCGGTAGGCGTGTGAGCGATGTGGCTCGGCT |
| ACTB | CGCAAATGGGCGGTAGGCGTGACA CAG AGTACTTGCGCTCAG |

**Supplementary Table 3.** *Primer sequences for qRT-PCR.*

| Gene | Forward Primer (5′-3′) | Reverse Primer (5′-3′) |
|---|---|---|
| ND1 | TCAAACTCAAACTACGCCCTG | GTTGTGATAAGGGTGGAGAGG |
| ND4 | CTCACACTCATTCTCAACCCC | TGTTTGTCGTAGGCAGATGG |
| ND5 | CTAGGCCTTCTTACGAGCC | CGCAAATGGGCGGTAGGCGTGT<br>TTGGGTTGAGGTGATGATG |
| ND6 | TGCTGTGGGTGAAAGAGTATG | CGCAAATGGGCGGTAGGCGTGC<br>CCATAATCATACAAAGCCCC |
| CO1 | GCCATAACCCAATACCAAACG | TTGAGGTTGCGGTCTGTTAG |
| CO2 | CTAGTCCTGTATGCCCTTTTCC | GTAAAGGATGCGTAGGGATGG |
| CO3 | CCTTTTACCACTCCAGCCTAG | CTCCTGATGCGAGTAATACGG |
| CYTB | CAATTATACCCTAGCCAACCCC | GGATAGTAATAGGGCAAGGACG |
| IRF7 | CTGTGGACACCTGTGACACC | TGCCCTCTCAGGAGCCAA |
| IFI27 | ATCAGCAGTGACCAGTGTGG | ATCAGCAGTGACCAGTGTGG |
| STAT1 | AACCTCGACAGTCTTGGCAC | CACTGAGACATCCTGCCACC |
| IFIT3 | GAAGGAACTGGGCCGCCTGCTAAG | GCCCTGGCCCATTTCCTCACTACC |
| IκBα | CTCCGAGACTTTCGAGGAAATAC | GCCATTGTAGTTGGTAGCCTTCA |
| MX1 | TTCTGGGTCGGAGGCTACAG | TGGATGGCGGCGTTCT |
| IRF3 | ACACATACTGGGCAGTGAGC | CTACAATGAAGGGCCCCAGG |
| IFI44 | CTGATTACAAAAGAAGACATGACAGAC | AGGCAAAACCAAAGACTCCA |
| IFITM1 | CGGCTCTGTGACAGTCTACC | TGCACAGTGGAGTGCAAAGG |
| ACTB | CCTGTACGCCAACACAGTGC | ATACTCCTGCTTGCTGATCC |
| GAPDH | CTCCTCCACCTTTGACGCTG | TCCTCTTGTGCTCTTGCTGG |

**Supplementary Table 4.** *Primer sequences for strand-specific qRT-PCR.*

| Gene | Forward primer (5′-3′) | Reverse primer (5′-3′) |
|---|---|---|
| ND1 Heavy | TCAAACTCAAACTACGCCCTG | CGCAAATGGGCGGTAGGCGTG |
| ND1 Light | GTTGTGATAAGGGTGGAGAGG | CGCAAATGGGCGGTAGGCGTG |
| ND4 Heavy | CTCACACTCATTCTCAACCCC | CGCAAATGGGCGGTAGGCGTG |
| ND4 Light | TGTTTGTCGTAGGCAGATGG | CGCAAATGGGCGGTAGGCGTG |
| ND 5 Heavy | CTAGGCCTTCTTACGAGCC | CGCAAATGGGCGGTAGGCGTG |
| ND5 Light | TAGGGAGAGCTGGGTTGTTT | CGCAAATGGGCGGTAGGCGTG |
| ND6 Heavy | TCATACTCTTTCACCCACAGC | CGCAAATGGGCGGTAGGCGTG |
| ND6 Light | TGCTGTGGGTGAAAGAGTATG | CGCAAATGGGCGGTAGGCGTG |
| CO1 Heavy | GCCATAACCCAATACCAAACG | CGCAAATGGGCGGTAGGCGTG |
| CO1 Light | TTGAGGTTGCGGTCTGTTAG | CGCAAATGGGCGGTAGGCGTG |
| CO2 Heavy | CTAGTCCTGTATGCCCTTTTCC | CGCAAATGGGCGGTAGGCGTG |
| CO2 Light | GTAAAGGATGCGTAGGGATGG | CGCAAATGGGCGGTAGGCGTG |
| CO3 Heavy | CCTTTTACCACTCCAGCCTAG | CGCAAATGGGCGGTAGGCGTG |
| CO3 Light | CTCCTGATGCGAGTAATACGG | CGCAAATGGGCGGTAGGCGTG |
| CYTB Heavy | CAATTATACCCTAGCCAACCCC | CGCAAATGGGCGGTAGGCGTG |
| CYTB Light | GGATAGTAATAGGGCAAGGACG | CGCAAATGGGCGGTAGGCGTG |
| GAPDH | CAACGACCACTTTGTCAAGC | CGCAAATGGGCGGTAGGCGTG |
| ACTB | ACACAGTGCTGTCTCGTGGTA | CGCAAATGGGCGGTAGGCGTG |

**Supplementary Table 5.** *Primer sequences for qRT-PCR in mouse samples.*

| Gene | Forward primer (5′-3′) | Reverse primer (5′-3′) |
|---|---|---|
| Nd1 | TCC GAG CAT CTT ATC CAC GC | GTA TGG TGG TAC TCC CGC TG |
| Nd4 | TAA TCG CAC ATG GCC TCA CA | CAT TTG AAG TCC TCG GGC CA |
| Nd5 | CAG CAC AAT TTG GCC TCC AC | TAG TCG TGA GGG GGT GGA AT |
| Nd6 | CCC GCA AAC AAA GAT CAC CC | TCT TGA TGG TTT GGG AGA TTG GT |
| Co1 | TCG GAG CCC CAG ATA TAG CA | TTT CCG GCT AGA GGT GGG TA |
| Co2 | CCT GGT GAA CTA CGA CTG CT | GGA CTG CTC ATG AGT GGA GG |
| Co3 | AAG GCC ACC ACA CTC CTA TTG | GCA GCC TCC TAG ATC ATG TGT |
| Cytb | TGC ATA CGC CAT TCT ACG CT | AGG CTT CGT TGC TTT GAG GT |
| Gapdh | CCC TTA AGA GGG ATG CTG CC | TAC GGC CAA ATC CGT TCA CA |
| Actb | GTA CTC TGT GTG GAT CGG TGG | AAC GCA GCT CAG TAA CAG TCC |

### Supplementary Materials and Methods

#### Patient saliva and tear samples

From 42 patients with SS, 31 patients with non-SS sicca, and 16 non-sicca voluntary donors, saliva (34 SS, 23 non-SS sicca, and 16 non-sicca controls) or tears (8 SS and 8 non-SS sicca samples) were collected. SS was classified according to the 2016 American College of Rheumatology/European League Against Rheumatism (ACR/EULAR) classification criteria for primary SS [1] and non-SS sicca subjects did not fulfill the ACR/EULAR criteria and the 2002 revised American-European Consensus Group criteria [2]. Their clinical characteristics are summarized in Table S1. Tear and saliva samples were stored at -80°C until further analysis.

#### RNA extraction and real-time quantitative polymerase chain reaction

To extract total RNAs from cultured cell, TRIzol (Ambion) was added directly to the cell pellet obtained after centrifugation at 10,000 *x g* for 30 sec. After precipitation of the extracted nucleic acids, DNase I (TaKaRa) was treated to remove DNA, and purified RNA was reverse transcribed using RevertAid reverse transcriptase (Thermo Fisher Scientific). For RNA extracted from patient samples and 3D spheroids, SuperScript IV Reverse Transcriptase (Invitrogen) was used to synthesize cDNA. For strand-specific RT-qPCR, reverse transcription primers containing CMV promoter sequences were designed to target the specific genes. cDNA was amplified by SYBR Green PCR master mix (Bioline) and RT-qPCR was performed using AriaMx Real-time PCR system (Agilent) and QuantStudio 1 Real-time PCR system (Thermo Fisher Scientific). Primers used in the study are provided in Table S2-S4. To analyze the localization of mtRNAs, cytosolic RNAs were isolated from cell lysates using the subcellular protein fractionation kit (Thermo Fisher Scientific) following the manufacturer's instruction. TRIzol LS (Ambion) was added individually at a 3:1 ratio to the cytosolic fraction to extract the RNAs. The rest of the steps were the same as above.

To extract RNA from NOD mice SG tissue, 20 μm-sections were dewaxed using xylene, then dehydrated in ethanol according to a previously published protocol [3]. Same reagents were used to synthesize cDNA and mice-specific primers were used for RT-qPCR analysis.

**Cell culture**

SV40-immortalized NS-SV-AC cell was provided by Professor Masayuki Azuma (Department of Oral Medicine, University of Tokushima Graduate Faculty of Dentistry, Japan) [4]. NS-SV-AC human salivary gland acinar cells were grown in Keratinocyte-SFM (Serum free media; Gibco) supplemented with 10% (v/v) heat-inactivated fetal bovine serum (FBS; Welgene) and 1% (v/v) 100x penicillin-streptomycin (Gibco). As for the primary SGEC obtained from SS patients, cells were cultured in Dulbecco's modified Eagle's Medium-Ham's F-12 (1:3) (Gibco) containing 2.5% fetal bovine serum (Gibco), 1% penicillin-streptomycin (Gibco), 0.4 μg/ml hydrocortisone (Sigma), 10 ng/ml epidermal growth factor (EGF; BioLegend), and 0.5 μg/ml insulin (Gibco). Single-cell suspensions were transferred to bovine type I collagen (PureCol; Advanced BioMatrix)-coated culture dishes. Patients were diagnosed as having primary SS according to the American-European Consensus Group criteria [2]. Informed consent was obtained from all patients according to the principles of the Declaration of Helsinki. Labial salivary gland biopsy tissues were obtained from control patients with sicca symptoms as well as from patients with primary SS. This study was approved by the Institutional Review Board of Seoul St. Mary's Hospital (approval no. KC13ONMI0646). Lip biopsy samples were minced in 1 unit/ml of Dispase solution (StemCell Technologies) containing 2 mg/ml of collagenase IV (Gibco) and were then digested with gentle resuspension. All cells were maintained at 37°C in a humidified 5% $CO_2$ incubator.

**Chemical treatment**

To induce dsRNA stress, 20 μg/ml of poly I:C (Sigma Aldrich) was transfected using Lipofectamine 3000 (Thermo Fisher Scientific) for 14 h following the manufacturer's guide. As for the controlled mock transfection, RNase-free water (Diethyl pyrocarbonate (DEPC)-treated water) was used.

To test the effect of Ach or SS-IgG on mt-dsRNAs induction, 100 μM of Ach (Sigma Aldrich) was co-treated with poly I:C transfection. Human IgG was prepared from 4 SS patients with anti-M3 muscarinic acetylcholine receptor antibodies [5], using the NAb$^{TM}$ Spin Kits for antibody purification (Thermo Fisher Scientific). IgG was pooled from four individual SS patients (125 μg/patient) and then mixed to the total mass of 500 μg, then lyophilized. The same method was used to pool IgG from four individual RA patients (125 μg/patient). IgG from human serum (Sigma Aldrich) was used as a control. All IgGs were dissolved in 150 mM NaCl before treatment. To examine the therapeutic effect of JAK1 inhibitor, 1 or 10 μg/mL of upadacitinib (MedKoo Biosciences) was treated 1 h prior to poly I:C transfection. To downregulate mt-dsRNAs expression, 20 μM of 2-CM (Santa Cruz Biotechnology) was pretreated 24 h prior to poly I:C transfection.

**Polymer thin films via the initiated chemical vapor deposition process**

A 300 nm-thick pV4D4 film was deposited directly onto tissue culture polystyrene (TCPS) via the initiated chemical vapor deposition (iCVD) process. TCPS was first placed on the stage of a custom-built iCVD chamber. V4D4 (97%; Jihyunchem Co.) was heated to 70°C for monomer vaporization. The vaporized monomer V4D4, and the initiator, *tert*-butyl peroxide (TBPO; 95%; Sigma Aldrich) were introduced into the iCVD chamber at flow rates of 1.94 and 0.744 sccm, respectively. The pressure of the chamber was set to 260 mTorr. To decompose TBPO to generate radicals, the filament temperature was heated to 200°C. During the deposition process, the stage was maintained to 38.5°C for the adsorption of monomers.

**RNA extraction and real-time quantitative polymerase chain reaction**

To extract total RNAs from cultured cell, TRIzol (Ambion) was added directly to the cell pellet obtained after centrifugation at 10,000 *x g* for 30 sec. After precipitation of the extracted nucleic acids, DNase I (TaKaRa) was treated to remove DNA, and purified RNA was reverse transcribed using RevertAid reverse transcriptase (Thermo Fisher Scientific). For RNA extracted from patient samples and 3D spheroids, SuperScript IV Reverse Transcriptase (Invitrogen) was used to synthesize cDNA. For strand-specific RT-qPCR, reverse transcription primers containing CMV promoter sequences were designed to target the specific genes. cDNA was amplified by SYBR Green PCR master mix (Bioline) and RT-qPCR was performed using AriaMx Real-time PCR system (Agilent) and QuantStudio 1 Real-time PCR system (Thermo Fisher Scientific). Primers used in the study are provided in Table S2-S4. To analyze the localization of mtRNAs, cytosolic RNAs were isolated from cell lysates using the subcellular protein fractionation kit (Thermo Fisher Scientific) following the manufacturer's instruction. TRIzol LS (Ambion) was added individually at a 3:1 ratio to the cytosolic fraction to extract the RNAs. The rest of the steps were the same as above.

To extract RNA from NOD mice SG tissue, 20 μm-sections were dewaxed using xylene, then dehydrated in ethanol according to a previously published protocol [3]. Same reagents were used to synthesize cDNA and mice-specific primers were used for RT-qPCR analysis.

**Western Blotting**

Cell lysates were prepared by incubating cells in the lysis buffer (50 mM Tris-HCl pH 8.0, 100 mM KCl, 0.5% NP-40, 10% Glycerol, and 1 mM DTT) supplemented with a protease inhibitor cocktail (Merck) followed by sonication. 30 - 40 μg of protein samples were loaded and separated on a 10% SDS-PAGE gel and transferred to a PVDF membrane (Merck) using an Amersham semi-dry transfer system. The primary antibodies used in this study were as follows: pPKR (Abcam, Ab81303), PKR (Cell Signaling Technology, D7F7), and β-tubulin (Cell Signaling Technology).

**Acid phosphatase assay (APH)**

Cells (or spheroids) were centrifuged for 10 min at 400 *x g* for spin down. After washing the pellets twice with DPBS (Welgene, LB001-02), the supernatant was discarded to obtain the final volume of 100 μl. 100 μl of APH assay buffer (0.1 M sodium acetate, 0.1% (v/v) Triton X-100 (Promega), 2 mg/ml Immunopure PNPP (Sigma Aldrich) in deionized/distilled water) was added to each well and incubated for 90 min at 37°C with 5% $CO_2$. After incubation, 10 μl of 1N NaOH was added, and absorption at 405 nm was measured on a microplate reader (BioTek). For each experiment, cell viability was normalized to the control group without Mock transfection.

### Immunocytochemistry

Spheroids from pV4D4-coated plates were transferred to a 1.5 ml tube, and analysis for 2D culture was performed on a confocal dish (SPL). Cells were fixed in 4% (w/v) paraformaldehyde for 15 min at room temperature. Fixed cells were then permeabilized with 0.3% (w/v) Triton X-100 (Sigma Aldrich) in Dulbecco's phosphate-buffered saline (DPBS) for 10 min at room temperature and blocked in 3% bovine serum albumin (BSA100; Bovogen) for 1 h. Cells were incubated with primary antibodies diluted in 1% BSA at a 1:100 ratio overnight at 4°C. The primary antibodies used in this study include: Occludin (Invitrogen; OC-3F10), ZO-1 (Cell Signaling Technology; 8193S), and AQP5 (Cell Signaling Technology; 59558S). Cells were then washed with DPBS and incubated with Alexa fluor 488-conjugated anti-mouse secondary antibody (Thermo Fisher Scientific; A31570) and Alexa fluor 555 conjugated anti-mouse secondary antibody (Thermo Fisher Scientific; A-21202) diluted at a 1:1000 ratio for 45 min at room temperature. DAPI was incubated for 5 min. All washing steps for spheroids include centrifugation (5 min, 200 g) and supernatant discarding. Fluorescent images were obtained using a confocal laser-scanning microscope (LSM 880, Carl Zeiss). The fluorescence intensity per area was quantified as integrated density divided by the area of each spheroid and then normalized to the control without poly I:C transfection.

### PKR fCLIP

To prepare PKR antibody-conjugated beads, protein A beads (Thermo Fisher Scientific) were incubated with PKR antibody (Cell Signaling Technology) in the fCLIP lysis buffer (20 mM Tris-HCl, pH 7.5, 15 mM NaCl, 10 mM EDTA, 0.5% NP-40, 0.1% Triton X-100, 0.1% SDS, and 0.1%

sodium deoxycholate) for 3 h at 4°C after adjusting NaCl concentration to 150 mM. Harvested cells were fixed with 0.1% (w/v) paraformaldehyde (Sigma Aldrich) for 10 min and immediately quenched by adjusting glycine (Bio-basic) concentration to 250 mM for an additional 10 min at room temperature. The cross-linked cells were lysed using the fCLIP lysis buffer for 10 min on ice and then sonicated. The NaCl concentration of the lysate was adjusted to 150 mM, and cell debris was separated by centrifugation. The lysate was added to the PKR antibody-conjugated beads and incubated for 3 h at 4°C. The beads were washed 4 times with the fCLIP wash buffer (20 mM Tris-HCl, pH 7.5, 150 mM NaCl, 10 mM EDTA, 0.1% NP-40, 0.1% SDS, 0.1% Triton X-100, and 0.1% sodium deoxycholate) and PKR-dsRNA complex was eluted from the beads by incubating in the elution buffer (200 mM Tris-HCl, pH 7.4, 100 mM NaCl, 20 mM EDTA, 2% SDS, and 7 M Urea) for 3 h at 25°C. The eluate was treated with 2 mg/ml proteinase K (Sigma Aldrich) for overnight at 65°C. RNA was purified using acid-phenol:Chloroform pH 4.5 (Thermo Fisher Scientific).

**Differentially expressed gene (DEG) analysis**

For DEG analysis, DESeq2 analysis (Love et al., 2014) was performed three times separately in each of DMSO and 2-CM-pretreated groups. Within each group, raw counts of two biological replicates with poly I:C transfection were compared against the raw counts of two replicates with the control group (i.e. DEPC transfection). The counts were normalized by DESeq2 during analysis. The analysis yielded three separate lists of differentially expressed genes, and the genes were ordered by the lowest adjusted p-values for further analysis.

**Supplementary Figure Legend**

**Supplementary Figure 1. Association of the five most significant mitochondrial RNA expression levels in body fluid with exocrine dysfunction.** (A) Top five ranked mitochondrial transcripts in tear or saliva samples in SS versus non-SS sicca patients. (B) ROC curve of the top five ranked RNAs in tear or saliva samples. 95% CI = 95% confidence interval. (C) Tear *CYTB* heavy strand RNA levels in SS versus non-SS sicca patients. (D) Negative correlation between tear *CYTB* heavy strand RNA levels and tear production (Schirmer test; p-value = 0.006 by Pearson's correlation, rho = -0.651). (E) Positive correlation between tear *CYTB* heavy strand RNA levels and the corneal and conjunctival damage (ocular surface staining score; p-value < 0.0001, rho = 0.811). (F) Upregulation of salivary *ND5* heavy RNA levels in SS patients (p-value < 0.0001 by one-way ANOVA with post hoc Tukey test). (G) Salivary *ND5* levels were not associated with unstimulated whole salivary flow rates. (H) Negative correlation between salivary ND5 levels and stimulated whole salivary flow rate (p-value = 0.01, rho = -0.343). (I) Positive correlation of *ND5* heavy strands with lymphocyte infiltration grades in the minor salivary glands (p-value = 0.001, rho = 0.494). Error bar means SEM and the dotted line represents the 95% confidence intervals. *** is p-values ≤ 0.001.

**Supplementary Figure 2. dsRNA stimulation leads to increased mt-dsRNA expression.** (A) elative mRNA expression of indicated ISGs in poly I:C stimulated cells. (B) Western blot analysis of PKR phosphorylation upon poly I:C stimulation. (C) APH assay showing cell viability upon poly I:C stimulation. (D) RNA levels of total mtRNAs and (E) strand-specific mtRNAs upon poly I:C stimulation. (F) Representative fluorescence images of *ND5* light and heavy strands upon poly I:C transfection obtained by RNA-FISH. The scale bars represent 50 μm. (G) Cytosolic export of mt-dsRNAs upon poly I:C transfection. (H) PKR-fCLIP-qPCR analysis showing enhanced interaction between mt-dsRNAs and PKR upon poly I:C stimulation. Unless mentioned otherwise, shown are three independent experiments with error bars denoting SEM. Cq values are relative to *ACTB* mRNA (unless mentioned otherwise), then normalized to Mock transfected control. All statistical significances were calculated using one-tailed Student's t-tests, *$p \leq 0.05$, **$p \leq 0.01$, and ***$p \leq 0.001$.

**Supplementary Figure 3. Poly I:C stimulation leads to mt-dsRNA induction in primary SGECs isolated from SS patients.** (A) Relative mt-dsRNA expression between primary SGECs of SS patients and non-SS sicca patients. (B) mt-dsRNA expression upon poly I:C stimulation in SGECs isolated from SS patients (patient #1 and #2) and (C) non-SS sicca patients (patient #3 and #4). Cq values are normalized to *GAPDH* mRNA, and then normalized to Mock transfected control. Shown are three independent experiments with error bars denoting SEM. All statistical significances were calculated using one-tailed Student's t-tests, *$p \leq 0.05$, **$p \leq 0.01$, and ***$p \leq 0.001$.

**Supplementary Figure 4. The 3D culture system enhances the expression of TJC and AQP5 proteins.** (A) Representative images of NS-SV-AC cell growth in 2D (TCPS) and 3D spheroids on a pV4D4-coated plate. (B) Representative images of NS-SV-AC spheroids on pV4D4-coated plates compared to conventional ULA 3D plates. The scale bars represent 200 μm. (C) Representative images of ZO-1 and Occludin expression in NS-SV-AC grown in 2D versus 3D. The scale bar represents 100 μm.

**Supplementary Figure 5. mRNA-seq data validation using qRT-PCR and GO analysis of 2-CM-pretreated samples.** (A) Relative mRNA expression of individual ISGs analyzed by qRT-PCR. All Cq values are relative to *ACTB* mRNA. Shown are three independent experiments with error bars denoting SEM. All statistical significances were calculated using one-tailed Student's t-tests, $*p \leq 0.05$, $**p \leq 0.01$, and $***p \leq 0.001$. (B) GO analysis of top 200 genes whose $\log_2$ fold change was rescued by 2-CM pretreatment upon poly I:C stimulation.

**Supplementary Figure 1.**

**A**

| mtRNA | AUC [95% CI] | P values |
|---|---|---|
| Tear | | |
| CYTB heavy | 1.00 [1.00-1.00] | 0.001 |
| ND4 heavy | 0.98 [0.94-1.00] | 0.001 |
| ND1 light | 0.97 [0.89-1.00] | 0.002 |
| CO1 heavy | 0.94 [0.81-1.00] | 0.003 |
| CO2 heavy | 0.94 [0.81-1.00] | 0.003 |
| CO2 light | 0.94 [0.81-1.00] | 0.003 |
| Saliva | | |
| ND5 heavy | 0.78 [0.66-0.90] | <0.0001 |
| CO1 heavy | 0.74 [0.61-0.87] | 0.002 |
| ND1 light | 0.73 [0.60-0.87] | 0.017 |
| ND6 light | 0.73 [0.59-0.87] | 0.004 |
| CYTB heavy | 0.72 [0.59-0.85] | 0.005 |

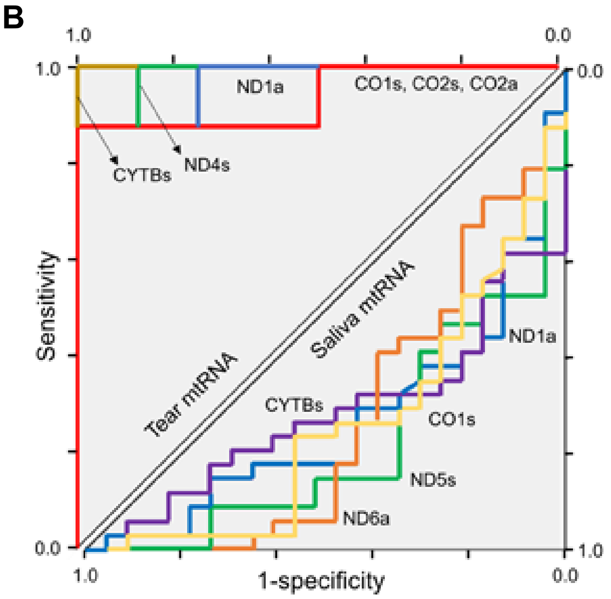


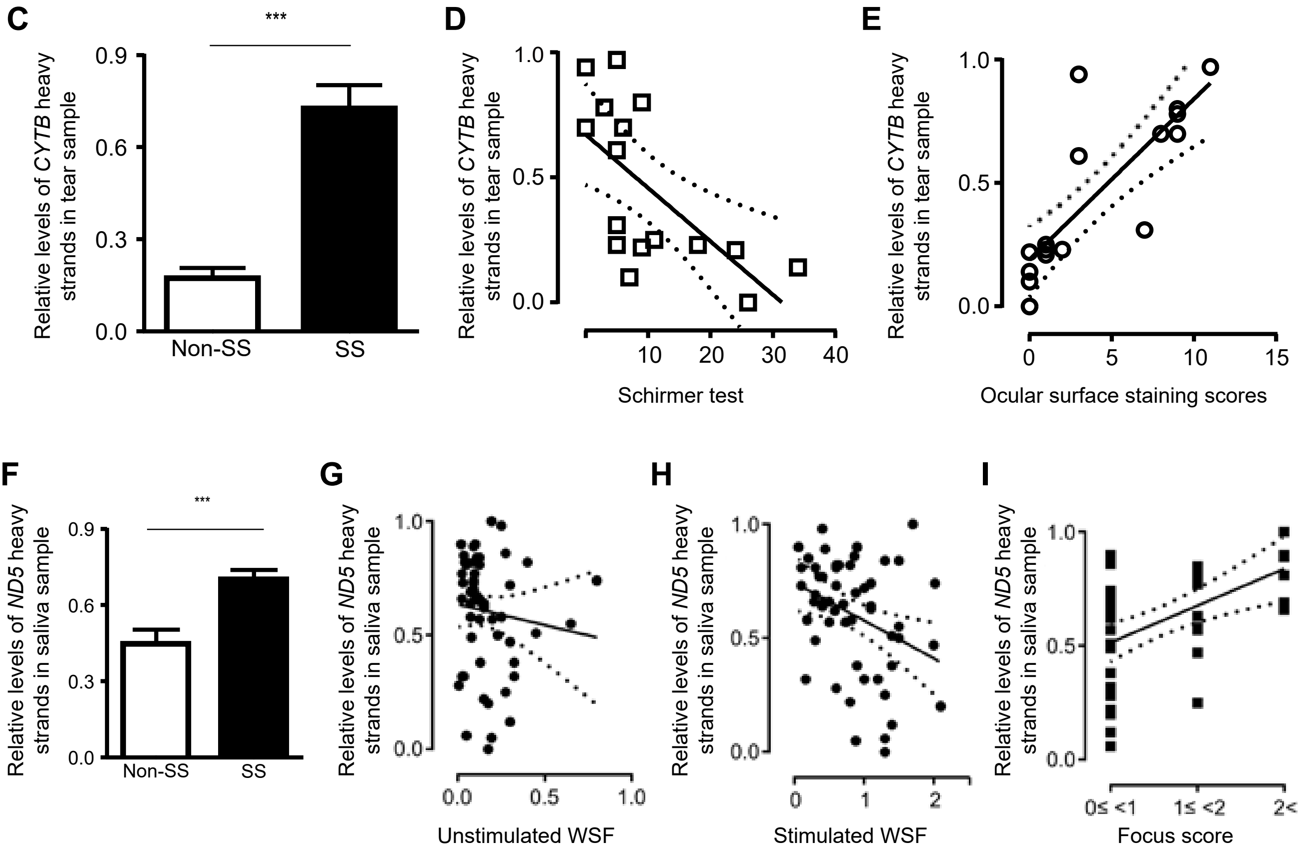

Supplementary Figure 2.

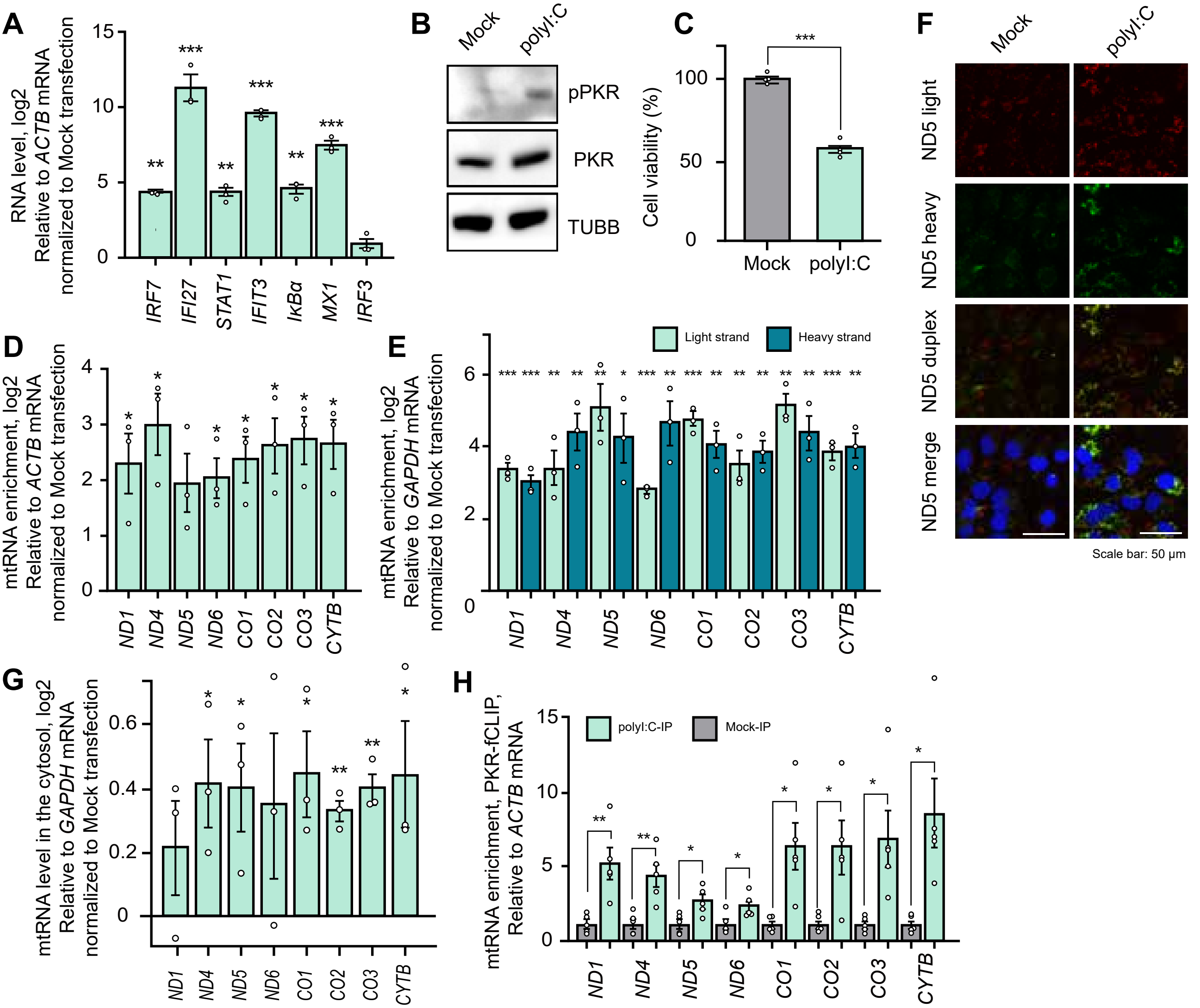

**Supplementary Figure 3.**

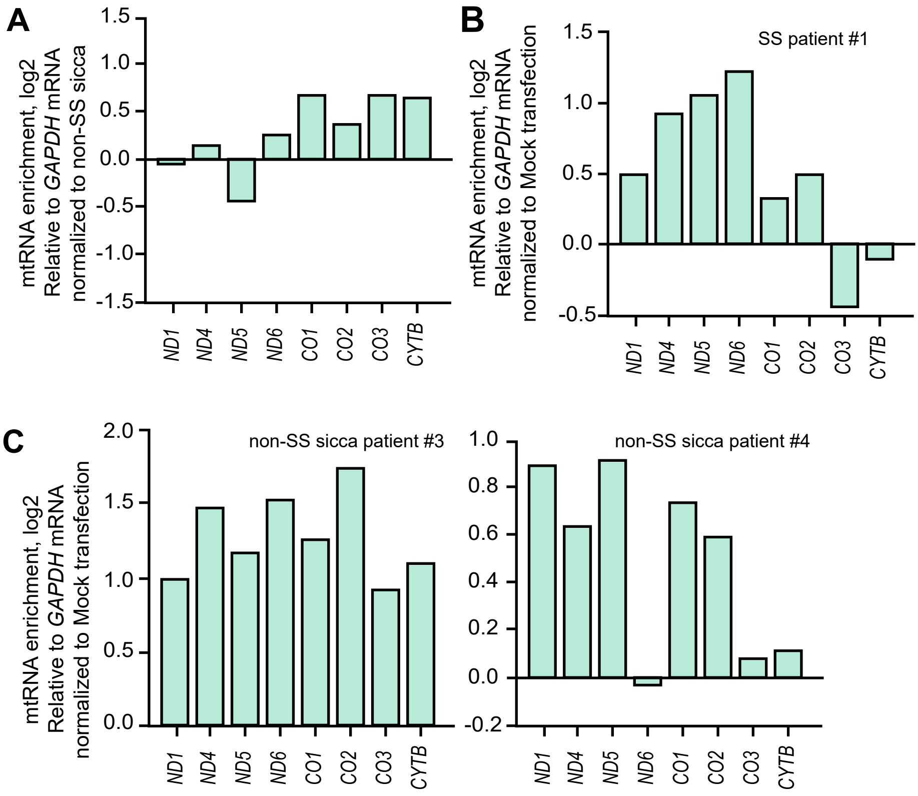

### Supplementary Figure 4.

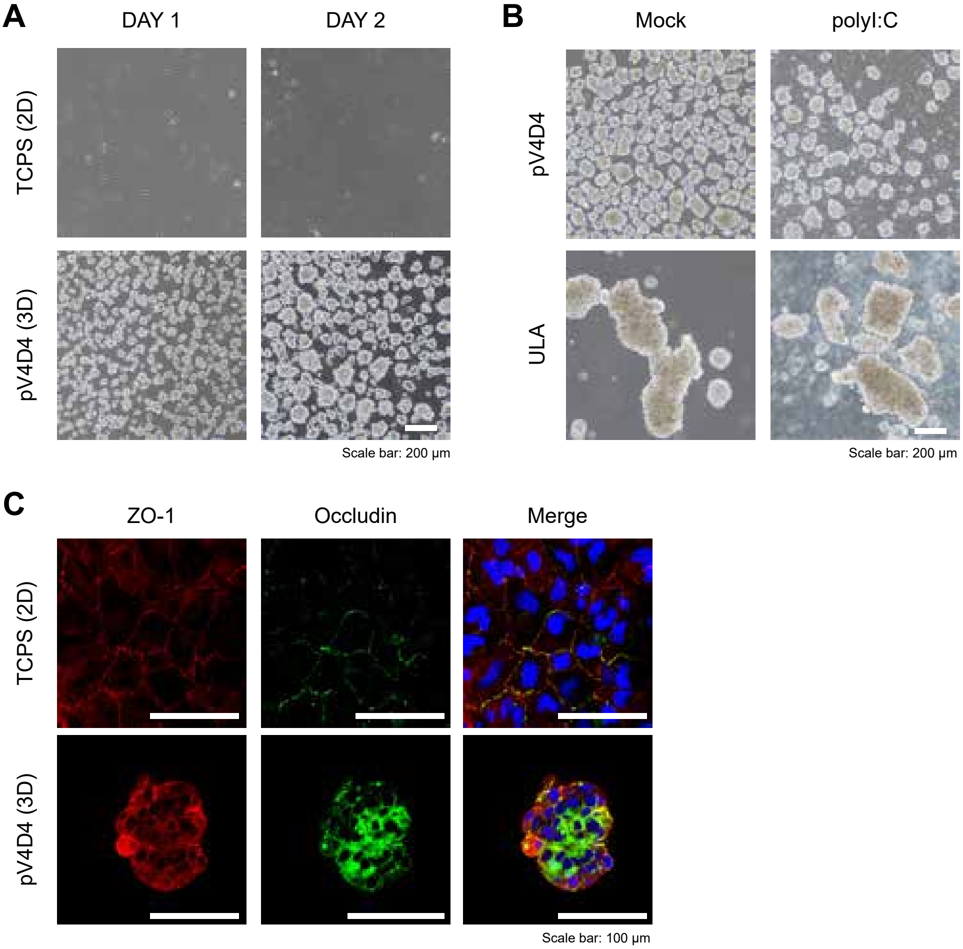

**Supplementary Figure 5.**

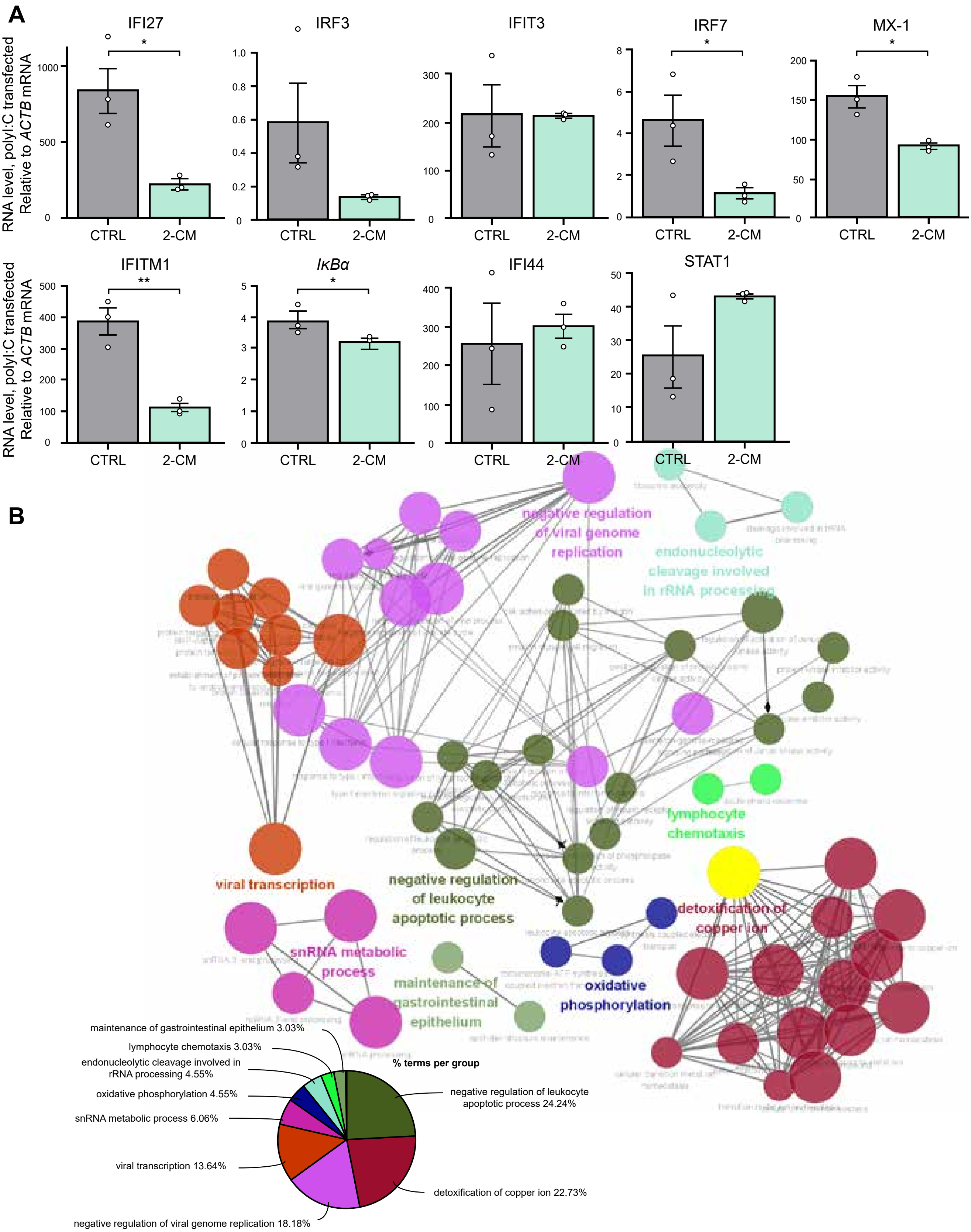
